# Unique Epigenetic Programming Distinguishes Regenerative Spermatogonial Stem Cells in the Developing Mouse Testis

**DOI:** 10.1101/674457

**Authors:** Keren Cheng, I-Chung Chen, Benjamin J. Hale, Brian P. Hermann, Christopher B. Geyer, Jon M. Oatley, John R. McCarrey

## Abstract

Spermatogonial stem cells (SSCs) both self-renew and give rise to progenitor spermatogonia that enter steady-state spermatogenesis in the mammalian testis. However, questions remain regarding the extent to which SSCs and progenitors represent stably distinct spermatogonial subtypes. Here we provide the first multiparametric integrative analysis of mammalian germ cell epigenomes comparable to that done by the ENCODE Project for >100 somatic cell types. Differentially expressed genes distinguishing SSCs and progenitors showed distinct histone modification patterns as well as differences in distal intergenic low-methylated regions. Motif-enrichment analysis predicted transcription factors that regulate this spermatogonial subtype-specific epigenetic programming, and gene-specific chromatin immunoprecipitation analyses confirmed subtype-specific differences in binding of a subset of these factors to target genes. Collectively, these results suggest that SSCs and progenitors are stably distinct spermatogonial subtypes differentially programmed to either self-renew and maintain regenerative capacity as SSCs, or lose regenerative capacity and initiate lineage commitment as progenitors.

## Introduction

An average adult human male produces 85-100 million sperm per day, all of which emanate from the highly proliferative seminiferous epithelium in the testis^1^. Within this epithelium spermatogenesis is sustained by spermatogonial stem cells (SSCs), daughters of which either replenish the SSC pool or contribute to the spermatogenic differentiation pathway as transit amplifying progenitors^2^. SSCs are a specialized subset of undifferentiated spermatogonia that can be functionally distinguished in the mouse model on the basis of a quantifiable transplantation assay^3, 4^ analogous to the transplantation assay reliably used for decades to identify hematopoietic stem cells^5^.

In the postnatal mouse testis, prospermatogonia give rise to undifferentiated spermatogonia of which only a subset become foundational SSCs^6, 7, 8^. The remaining undifferentiated spermatogonia become progenitors primed to initiate spermatogenic differentiation ^9^ or undergo cell death^10, 11, 12, 13, 14^. Conflicting theories describe the dynamics by which mammalian SSCs acquire their fate in the developing testis and/or maintain their fate in the adult testis. The long-standing “A_single_ model” (A_s_ model) holds that individual A_s_ spermatogonia represent SSCs that divide to either self-renew or give rise to paired (A_pr_) and then aligned chains (A_al-4-16_) of progenitor spermatogonia connected by intercellular bridges^15, 16^. The “revised A_s_ model” suggests that a subset of A_s_ spermatogonia – distinguishable by expression of high levels of a marker transgene (*Id4-eGfp*) – function as self-renewing SSCs, while the remaining A_s_ spermatogonia represent a transient subpopulation en route to becoming progenitors^11^. The “fragmentation model” suggests that SSC fate can be adopted or lost by individual spermatogonia via transition between the A_s_ and various A_pr_-A_al-4-16_ states, <u>or</u> vice versa^17^. The A_s_ model (original or revised) predicts that SSCs are fundamentally distinct from progenitors. The fragmentation model, on the other hand, holds that all undifferentiated spermatogonial subtypes are equipotent, such that A_s_ spermatogonia can either self-renew or give rise to A_pr_-A_al-4-16_ spermatogonia that can, in turn, either continue spermatogenic differentiation <u>or</u> revert back to the A_s_ subtype^17^.

Studies based on detection of specific marker proteins^18, 19^, lineage tracing^17^, or bulk^20^ or single-cell^21, 22, 23^ RNA-sequencing (RNA-seq), have confirmed that undifferentiated spermatogonia display heterogeneous patterns of gene expression, indicating distinct spermatogonial subpopulations including SSCs, progenitors, transitory cells and cells undergoing cell death^9, 10, 11, 24, 25, 26^. Expression of the *Id4-eGfp* transgene marks a majority of undifferentiated spermatogonia, including transplantable/regenerative SSCs^27^. Selective FACS-based recovery of the brightest (ID4-eGFP^Bright^) and dimmest (ID4-eGFP^Dim^) portions of ID4-eGFP+ spermatogonia significantly enriches regenerative SSCs or non-regenerative progenitors, respectively^20, 27^. Similarly, dual FACS-based selection of ID4-EGFP+ cells expressing high or low levels of the endogenous cell surface marker, TSPAN8, significantly enriches or depletes transplantable SSCs^28^. These subpopulations of SSC-enriched or progenitor-enriched spermatogonia express DEGs encoding factors favoring self-renewal and maintenance of a stem cell state, versus proliferation and commitment to spermatogenic differentiation, respectively^20, 21, 28^.

We reasoned that if SSCs and progenitors represent fundamentally distinct spermatogonial subtypes, distinguishable epigenomic programming profiles should be associated specifically with DEGs. Indeed, the ENCODE and Roadmap Epigenomics and related projects reported distinct transcriptomes accompanied by up to 15 unique cell-type specific epigenetic programming profiles at promoters and enhancers for more than 100 different somatic cell types in mammals^29, 30, 31, 32, 33, 34, 35, 36^. However, none of these studies examined germ cells. Here, we used FACS to selectively recover highly enriched subpopulations of ID4-eGFP^Bright^ regenerative SSCs (“SSC-enriched spermatogonia”) and ID4-eGFP^Dim^ non-regenerative progenitors (“progenitor-enriched spermatogonia”) to perform multi-parametric integrative analysis of genome-wide patterns of DNA methylation, six different histone modifications, and chromatin accessibility in conjunction with subtype-specific transcriptome analysis to identify unique epigenetic landscapes associated with DEGs. We then performed motif enrichment analysis followed by indirect immunofluorescence (IIF) and chromatin immunoprecipitation (ChIP) to identify candidate factors that may either direct establishment, or mediate effects of differential epigenetic programming of spermatogonial-subtype specific genes. Our results provide unprecedented insight into the epigenetic programming associated with DEG patterns that distinguish SSCs and progenitors, and suggest that SSCs represent a unique spermatogonial subtype epigenetically programmed to retain SSC function, whereas progenitors have transitioned to a distinct fate associated with lineage commitment and spermatogenic differentiation.

## Results

Quadruplicate samples of regenerative SSC-enriched and non-regenerative progenitor-enriched spermatogonia were selectively recovered from testes of postnatal day 6 (P6) *Id4-eGfp* transgenic mice by FACS sorting for relative eGFP fluorescence as previously described^20^. Each epigenomic assay was run on four different samples of ID4-eGFP^Bright^ and ID4-eGFP^Dim^ cells to assess genome-wide patterns of DNA methylation, six different histone modifications, chromatin accessibility and gene expression. Each assay was conducted on identical aliquots of each sample, rendering results of each directly comparable.

### Differential Gene Expression Distinguishes Regenerative SSC-Enriched and -Depleted Spermatogonial Subpopulations

Previous bulk and single-cell RNA-seq (scRNA-seq) analyses of SSC- and progenitor-enriched spermatogonial subpopulations in the developing testis^20,21^ revealed distinct patterns of differential gene expression. Here, we first conducted bulk RNA-seq as a context for our bulk epigenomics analyses (Fig. 1a), and then used results from our previous scRNA-seq analysis^21^ to further delineate spermatogonial-subtype specific gene expression patterns (Fig. 1b). We identified nine distinct cellular subtype clusters, of which six were spermatogonial subtypes (Fig. 1b, clusters 1-4,6,7) and three were somatic cell types (Fig. 1b, clusters 5,8,9) based on expression of known cell-type specific marker genes (Fig. 1c). Spermatogonial clusters resolved into two subsets – those representing predominantly ID4-eGFP^Bright^ cells (Fig. 1b, clusters 1,3,4,7) and those representing predominantly ID4-eGFP^Dim^ cells (Fig. 1b, clusters 2,6), exemplifying the consistency of differential gene expression distinguishing these two spermatogonial subpopulations^21^. 1211 genes found to be highly expressed in somatic cell clusters in our scRNA-seq data were subsequently excluded from our bulk RNA-seq datasets that were then used for all subsequent comparisons with our bulk epigenomics datasets. This refined bulk RNA-seq data revealed 21,234 genes expressed in either ID4-eGFP^Bright^ or ID4-eGFP^Dim^ spermatogonia, or both. Of these, 669 genes were up-regulated [log_10_-fold difference of ≥1.5x (*p* < 0.01)] in ID4-eGFP^Bright^ spermatogonia (= “Class 1 genes”), 373 were up-regulated in ID4-eGFP^Dim^ spermatogonia (= “Class 2 genes”), and 20,192 were expressed at similar levels in both subpopulations (= “Class 3 genes”), including examples shown in Figure 1d and Table S1.

**Fig. 1:**
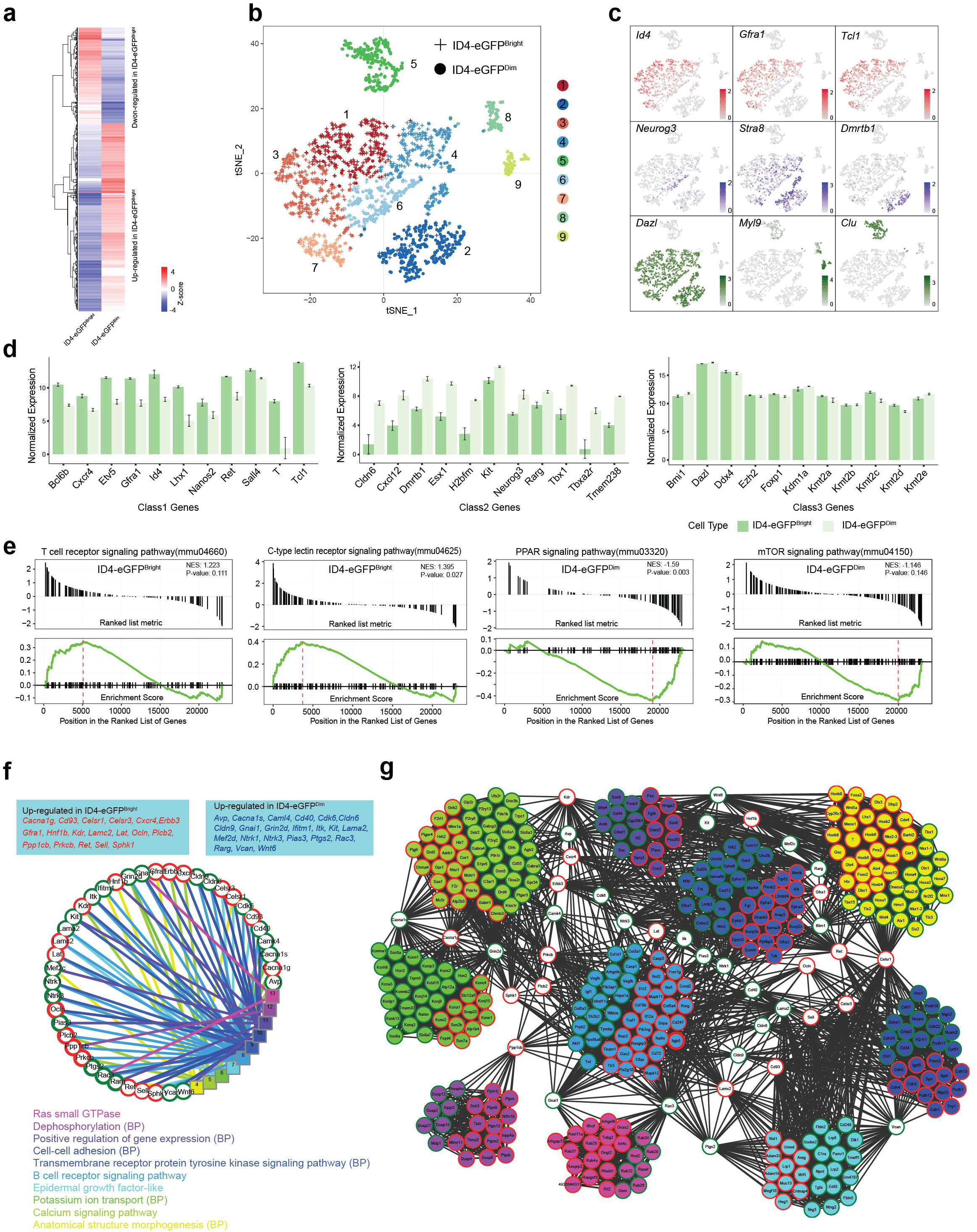
Differential gene expression in SSC-enriched and progenitor-enriched spermatogonial subpopulations. **a** Differential gene expression profiling of ID4-eGFP^Bright^ and ID4-eGFP^Dim^ spermatogonia by bulk RNA-seq. Genes with > 1.5 LFC (Log2 Fold Change difference > 1.5, p < 0.01) were hierarchically clustered. Red and blue colors indicate high and low expression levels in z-score. **b** Single cell RNA-seq (scRNA-seq) of P6 ID4-eGFP^Bright^ and ID4-eGFP^Dim^ spermatogonia. Each dot represents a single cell, + indicates an ID4-eGFP^Bright^ cell, indicates an ID4-eGFP^Dim^ cell. Nine distinct cell clusters were identified by Seurat. **c** scRNA-seq data showing expression of individual cell-type specific markers for SSCs (*Id4, Gfra1, Tcl1*), progenitors (*Neurog3, Stra8, Dmrtb1*) and early differentiated spermatogonia (*Dazl, Myl9, Clu*). **d** Normalized expression of genes significantly up-regulated in ID4-eGFP^Bright^ cells (Class 1 genes), up-regulated in ID4-eGFP^Dim^ cells (Class 2 genes), or constitutively expressed in both ID4-eGFP^Bright^ and ID4-eGFP^Dim^ cells (Class 3 genes). Data are presented as mean Å} SD and p < 0.01. **e** Gene Set Enrichment Analysis (GSEA) of KEGG pathways detected among genes differentially expressed in ID4-eGFP^Bright^ and ID4-eGFP^Dim^ spermatogonia. The T-cell receptor signaling and C-type lectin receptor signaling pathways are upregulated in ID4-eGFP^Bright^ spermatogonia, while the PPAR signaling and mTOR signaling pathways are up-regulated in ID4-eGFP^Dim^ spermatogonia. **f** Functional network analysis of genes differentially expressed in ID4-eGFP^Bright^ and ID4-eGFP^Dim^ spermatogonia. Red node color indicates up-regulated gene expression in ID4-eGFP^Bright^ cells, whereas green node color indicates up-regulated gene expression in ID4-eGFP^Dim^ cells. **g** Functional clustering of genes differentially expressed in ID4-eGFP^Bright^ and ID4-eGFP^Dim^ spermatogonia. Fill colors coordinate with those used in b to indicate distinct gene clusters, red outlines indicate genes up-regulated in ID4-eGFP^Bright^ cells, indicate genes upregulated in ID4-eGFP^Dim^ cells.

Gene sets enrichment of Kyoto Encyclopedia of Genes and Genomes (KEGG) analysis revealed pathways differentially enriched in SSC- and progenitor-enriched spermatogonia. The four most differentially enriched pathways included two that were up-regulated in ID4-eGFP^Bright^ and two that were up-regulated in ID4-eGFP^Dim^ spermatogonia (Fig. 1e). Elevated gene sets enrichment of the mTOR signaling pathway in progenitors was previously described ^37^, but that of the T cell receptor signaling pathway and the PPAR signaling pathway in SSCs and of the C-type lectin receptor pathway in progenitors is novel. Other enriched pathways, such as the RAP1 signaling pathway and the P13K-AKT signaling pathway, were expressed at similar levels in both ID4-eGFP^Bright^ and ID4-eGFP^Dim^ spermatogonia^38^ (Fig. S1; Table S1). Finally, functional gene networks (*FGNet*) analysis^39^ identified 10 functional metagroups among the DEGs (Fig. 1f) and 40 differentially expressed node or hub genes that interconnected these metagroups (Fig. 1g). Three of the metagroups included predominantly Class 1 genes, another was composed primarily of Class 2 genes, and six others were made up of similar proportions of Class 1 and Class 2 genes. In several cases, distinct sets of genes involved in similar functional groups were expressed in each spermatogonial subtype, and these appeared to be regulated by distinct hub genes. Thus, 18 hub genes were up-regulated in SSC-enriched spermatogonia, including *Erbb3, Gfra1*, and *Ret*, while 22 were up-regulated in progenitor-enriched spermatogonia, including *Kit, Rarg,* and *Wnt6* (Fig. 1f).

### Genic Region Patterns of Chromatin Modifications Are Associated with Gene Expression in SSC-Enriched and Progenitor-Enriched Spermatogonia

We analyzed aliquots of the same samples of each spermatogonial subtype by 1) ChIP-seq to detect six different histone modifications – H3K4me1,2,3, H3K9me3, H3K27ac, and H3K27me3, 2) ATAC-seq to assess chromatin accessibility, and 3) MeDIP-seq to examine DNA methylation and matched these results with our corresponding bulk RNA-seq data. K-means clustering of genic region data revealed six different patterns of histone modifications (Fig. 2a). Four modifications (H3K4me1,2,3 & H3K27ac) were enriched in genes expressed in one or both spermatogonial subpopulations, predominantly in promoter regions (Fig. 2a, clusters 1,2,3,5). Genes that were either not expressed or expressed at very low levels in one or both spermatogonial subpopulations (cluster 6) showed enrichment of the inactive H3K27me3 modification and depletion of the active H3K27ac modification (Fig. 2a). Genes which were not expressed in either subpopulation (cluster 4), showed enrichment of H3K4me1,2,3 and H3K27me3 within transcribed or downstream genic regions, but not at promoter regions. Within each cluster, enrichment of H3K4me1,2,3 and H3K27ac correlated positively with enhanced chromatin accessibility, and negatively with DNA methylation.

**Fig. 2:**
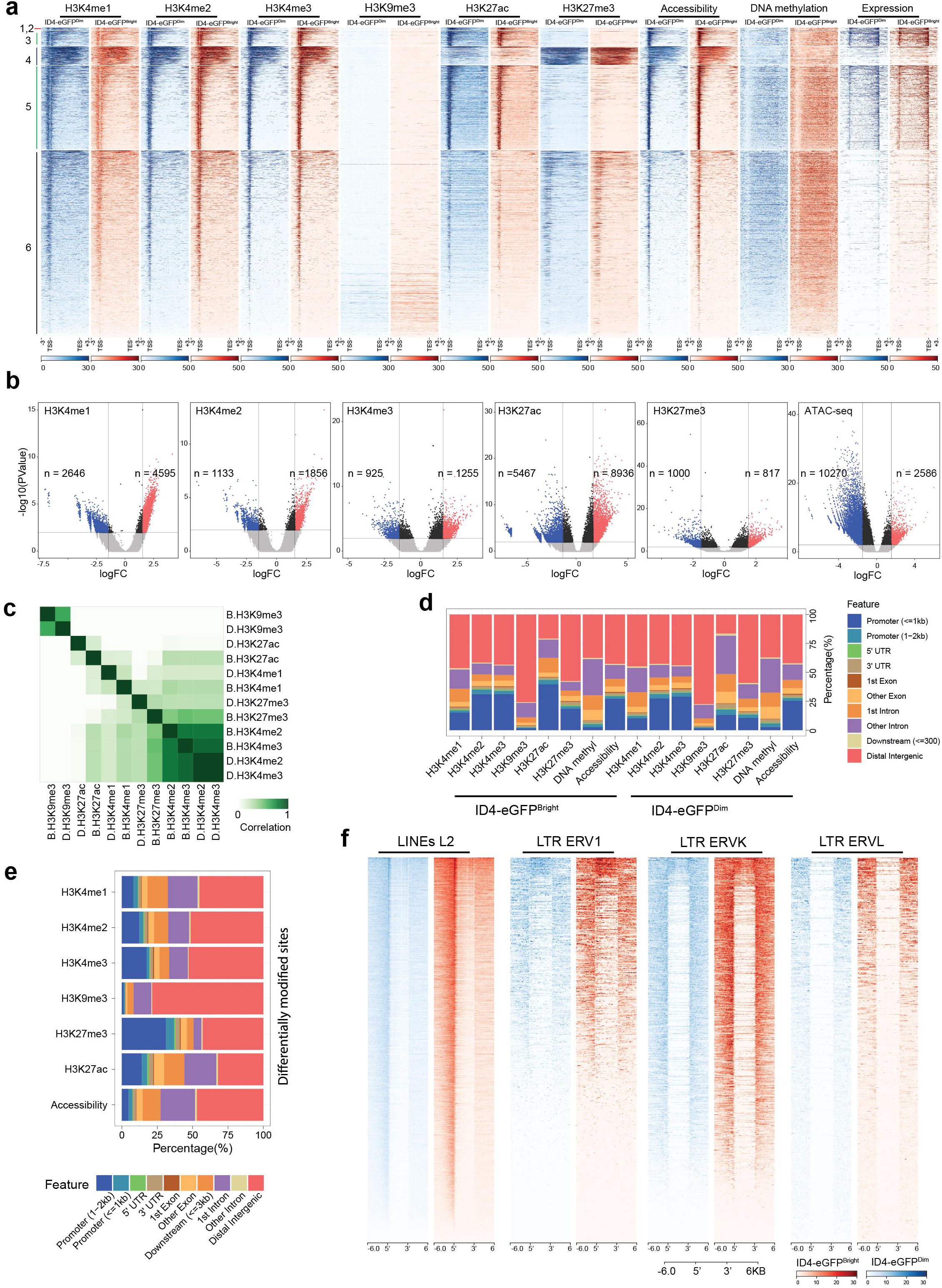
Epigenetic profiling of ID4-eGFP^Bright^ and ID4-eGFP^Dim^ spermatogonia. **a** Heatmaps show patterns of histone modifications (H3K4me1, H3K4me2, H3K4me3, H3K9me1, H3K27ac & H3K27me3) deduced by ChIP-seq, chromatin accessibility deduced by ATAC-seq, DNA methylation deduced by MeDIP-seq, and transcript abundance deduced by bulk RNA-seq within genic sequences of all mouse Refseq annotated genes (n = 24,012) in ID4-eGFP^Bright^ (red tracings) and ID4-eGFP^Dim^ (blue tracings) spermatogonia. All assays were conducted on aliquots of the same spermatogonial subtype samples. Sites are ordered by k-means clustering of signals between TSSs and TTSs (+ 3kb of upstream and 3kb of downstream sequence in each case). Blue color indicates reads from ID4-eGFP^Dim^ cells, red color indicates reads from ID4-eGFP^Bright^ cells. **b** Site-specific differences in enrichment of histone modifications and chromatin accessibility distinguishing mouse ID4-eGFP^Bright^ and ID4-eGFP^Dim^ spermatogonia. Red dots denote significantly enriched reads in ID4-eGFP^Bright^ cells and blue dots denote significantly enriched reads in ID4-eGFP^Dim^ cells for each modification or chromatin accessibility site. Black dots denote reads below LFC 1.5; grey dots denote p > 0.05. **c** Correlations among individual histone modification patterns in ID4-eGFP^Bright^ and ID4-eGFP^Dim^ spermatogonia. **d** Relative distribution of each histone modification as a function of genomic features genome wide in ID4-eGFP^Bright^ and ID4-eGFP^Dim^ spermatogonia. **e** Relative distribution of each type of histone modification showing differential enrichment in ID4-eGFP^Bright^ and ID4-eGFP^Dim^ spermatogonia as a function of genomic features genome-wide. **f** Heatmaps of H3K9me3 deposition in four types of repeats – LINEs(L2) and LTRs (*ERV1, ERVK, ERVL*) in ID4-eGFP^Bright^ and ID4-eGFP^Dim^ spermatogonia.

Significant (*p* < 0.05) differences in enrichment of each modification and of accessible chromatin were found in ID4-eGFP^Bright^ and ID4-eGFP^Dim^ spermatogonia (Fig. 2b). More accessible genomic regions were found in ID4-eGFP^Dim^ spermatogonia than in ID4-eGFP^Bright^ spermatogonia genome-wide (Fig. 2b). Occurrence of H3K4me2 and H3K4me3 modifications was more highly correlated than other pairs of modifications (Fig. 2c), and the prevalence of histone modifications was greatest in distal intergenic regions and promoters (Figs. 2d,e). Enrichment of the H3K9me3 modification occurred most prevalently in 5’ and 3’ regions of repeat elements^40^ (Fig. 2f). A GO analysis of gene promoters enriched for either H3K27ac or H3K27me3 was consistent with differential gene expression favoring enhanced maintenance of the stem cell state in SSC-enriched spermatogonia and of lineage priming in progenitor-enriched spermatogonia (Fig. S2).

### Epigenetic Landscapes Distinguish Promoters of Genes Differentially Expressed in SSC-Enriched and Progenitor-Enriched Spermatogonia

Exemplary sets of Class 1 genes up-regulated in SSC-enriched spermatogonia (*Tspan8, Pax7, Lhx1, Egr2, Gfra1, Id4, Tcl1*) and Class 2 genes up-regulated in progenitor-enriched spermatogonia (*Dmrtb1, Neurog3, Rarg, Kit, Lmo1, Crabp1*) illustrate differences in promoter programming associated with spermatogonial subtype-specific gene expression (Fig. 3a-h). Promoters of up-regulated genes showed enrichment of H3K4me3 and H3K27ac plus increased chromatin accessibility, while down-regulated genes showed decreased prevalence of H3K4me3 coupled with enrichment of H3K27me3 and decreased chromatin accessibility (Fig. 3a-h). Enrichment of H3K4me1,2,3 relative to that of H3K27me3 coincided with elevated expression in both spermatogonial subtypes (Fig. 3i,j). Collectively, our assessments revealed that, in each respective spermatogonial subpopulation, promoters of up-regulated genes showed: 1) enriched H3K4me1,2,3; 2) enriched H3K27ac; 3) depleted H3K27me3; 4) enhanced chromatin accessibility; and 5) hypomethylated DNA. In contrast, promoters of down-regulated genes showed: 1) decreased H3K4me1,2,3; 2) depleted H3K27ac; 3) enriched H3K27me3; 4) decreased chromatin accessibility; and 5) hypomethylated DNA. Thus, differential enrichment of H3K27ac or H3K27me3 in promoter regions correlated most closely with regulation of DEGs distinguishing ID4-eGFP^Bright^ and ID4-eGFP^Dim^ spermatogonia (Fig. 3k), as corroborated by heatmap and genome browser track data (Fig. S3).

**Fig. 3:**
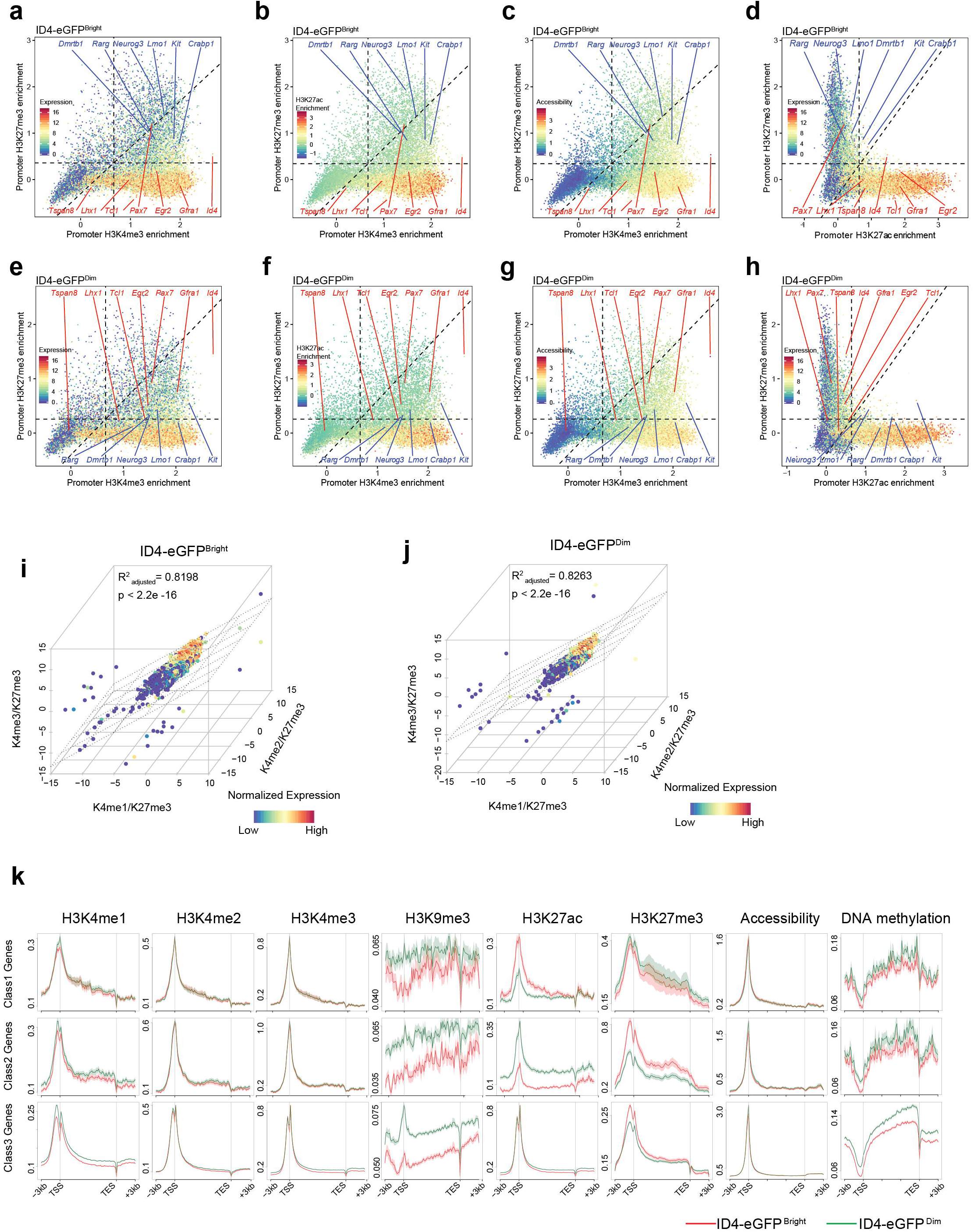
Epigenetic profiling at gene promoters in ID4-eGFP^Bright^ and ID4-eGFP^Dim^ spermatogonia. **a-h** Relative expression of genes as a function of enrichment of H3K27me3 and H3K4me3 in a ID4-eGFP^Bright^ (a) and ID4-eGFP^Dim^ (e) spermatogonia; relative enrichment of the H3K27ac modification as a function of enrichment of H3K27me3 and H3K4me3 in ID4-eGFP^Bright^ (b) and ID4-eGFP^Dim^ (f) spermatogonia; relative chromatin accessibility as a function of enrichment of H3K27me3 and H3K4me3 in ID4-eGFP^Bright^ (c) and ID4-eGFP^Dim^ (g) spermatogonia; and relative expression of genes as a function of enrichment of H3K27me3 and H3Kf27ac in ID4-eGFP^Bright^ (d) and ID4-eGFP^Dim^ (h) spermatogonia. Dashed lines indicate the minimum threshold value indicative of enrichment within each bimodal distribution. Each dot = 1 gene. Red dots = Class1 genes, blue dots = Class 2 genes. Exemplary known marker genes are identified. **i-j** Relative gene expression levels as a function of ratios of H3K4me3/H3K27me3, H3K4me1/H3K27me3 and H3K4me2/H3K27me3 in ID4-eGFP^Bright^ (i) and ID4-eGFP^Dim^ (j) spermatogonia. **k** Patterns of enrichment of individual histone modifications, chromatin accessibility and DNA methyhlation within genic regions of differentially expressed (Class 1 & Class 2) genes and similarly expressed (Class 3) genes in ID4-eGFP^Bright^ (red tracings) (i) and ID4-eGFP^Dim^ (green tracings) (j) spermatogonia. TSS = transcription start site, TES = transcription end site.

### Intergenic Enhancers Are Differentially Programmed in SSC-Enriched and Progenitor-Enriched Spermatogonia

Nearly 50% of accessible chromatin regions resided in distal intergenic regions in both ID4-eGFP^Bright^ and ID4-eGFP^Dim^ spermatogonia (Fig. 2d), and nearly all intergenic ATAC-seq peaks co-localized with peaks of H3K4me1,2,3 enrichment and hypomethylated DNA (Fig. 4a), indicative of enhancers ^41^. Epigenetic programming of enhancers is generally more variable than that of promoters ^42^ The three forms of H3K4me (1,2,3) typically co-located in both ID4-eGFP^Bright^ and ID4-eGFP^Dim^ spermatogonia (Fig. 4b,d), whereas enrichment of either H3K27ac (active enhancers), H3K27me3 (inactive or repressive enhancers), or chromatin accessibility + hypomethylated DNA without enrichment of H3K27ac or H3K27me3 (primed enhancers)^41^ appeared as distinct, non-colocalized modifications (Fig. 4c,e). These data revealed 5,546 active, 9,055 inactive, and 8,878 primed enhancers similarly programmed in both spermatogonial subtypes (Fig. 4f-h). However, an additional 3,024 active enhancers, 3,220 inactive enhancers and 5,952 primed enhancers were found uniquely in ID4-eGFP^Bright^ cells, while 2,979 active enhancers, 3,117 inactive enhancers and 7,660 primed enhancers were unique to ID4-eGFP^Dim^ cells. GO terms of genes associated with enhancers uniquely active, inactive or primed in SSC-enriched spermatogonia versus those in progenitor-enriched spermatogonia were consistent with maintenance of an undifferentiated stem cell state in SSCs and cell fate priming and differentiation in progenitors (Fig. 4i-k). As expected, genes associated with active enhancers in each spermatogonial subtype were up-regulated in that subtype (Fig. S4).

**Fig. 4:**
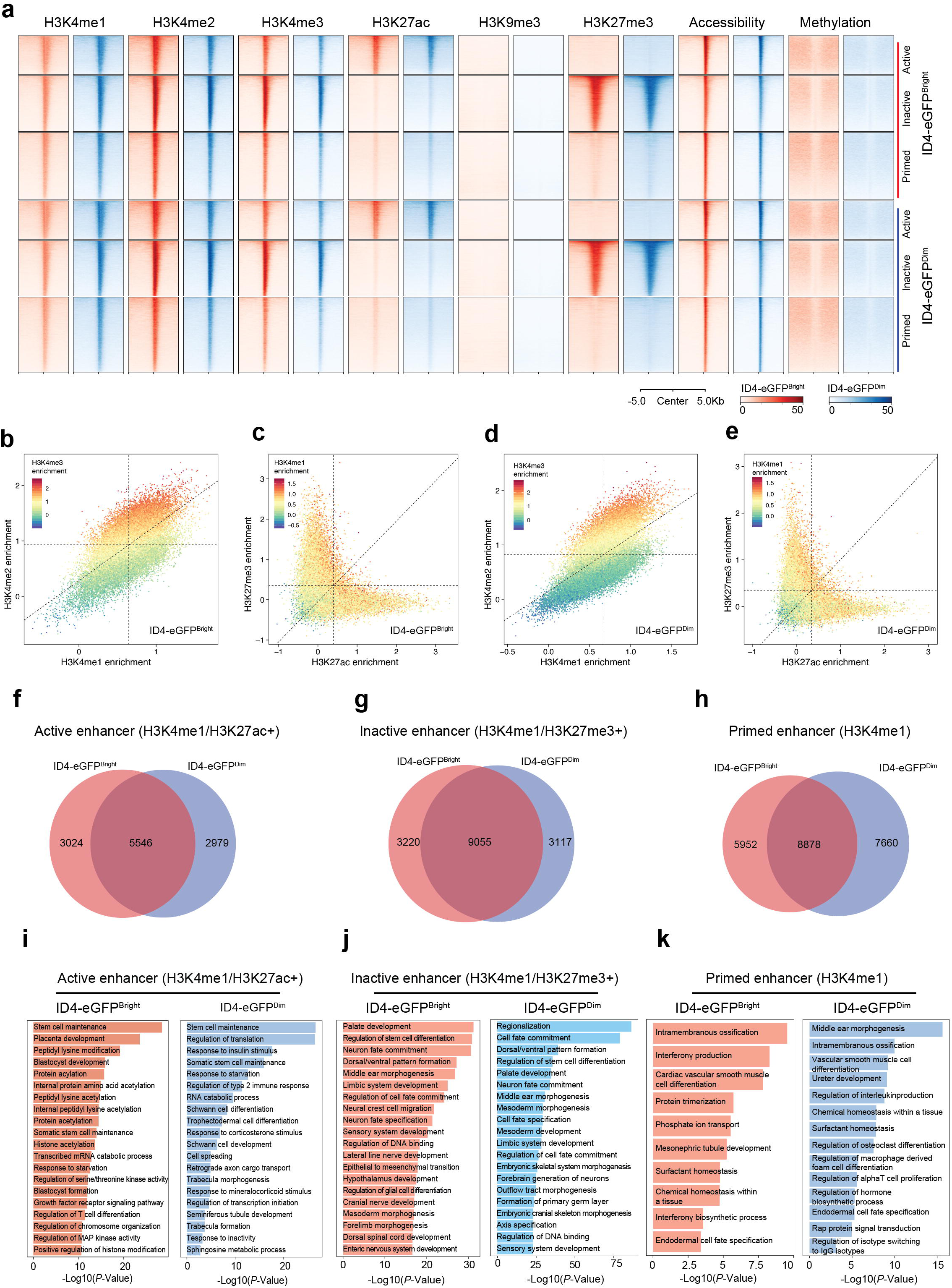
Epigenetic profiling at enhancers in ID4-eGFP^Bright^ and ID4-eGFP^Dim^ spermatogonia. **a** Heatmaps show patterns of enrichment of each histone modification, chromatin accessibility and DNA methylation at sites of intergenic enhancers in ID4-eGFP^Bright^ (red tracings) and ID4-eGFP^Dim^ (blue tracings) spermatogonia. **b-e** Dot plots showing positive or negative combinatorial enrichment of different histone modifications at enhancers. Dashed lines indicate the minimum threshold value indicative of enrichment within each bimodal distribution. Each dot = 1 enhancer. **b** Enrichment of the H3K4me3 modification correlates positively with enrichment of the H3K4me2 and H3K4me1 modifications. **c** Enrichment of the H3K4me1 modification correlates positively with enrichment of either the H3K27me3 or the H3K27ac modifications, but enrichment of the H3K27me3 modification correlates negatively with enrichment of the H3K27ac modification, and vice versa. **d** Enrichment of the H3K4me3 modification correlates positively with enrichment of either the H3K27me3 or the H3K27ac modifications, but enrichment of the H3K27me3 modification correlates negatively with enrichment of the H3K27ac modification, and vice versa. **f-h** Venn diagrams showing proportions of active (f), inactive (g) and primed (h) enhancers unique to ID4-eGFP^Bright^ spermatogonia, common to both ID4-eGFP^Bright^ and ID4-eGFP^Dim^ spermatogonia, or unique to ID4-eGFP^Dim^ spermatogonia. **i-k** GREAT GO analysis of functions encoded by genes associated with active (i), inactive (j) or primed (k) enhancers in ID4-eGFP^Bright^ or ID4-eGFP^Dim^ spermatogonia.

### Differentially Methylated Regions in SSC-Enriched and Progenitor-Enriched Spermatogonial Subpopulations Are Located Primarily in Intergenic Regions

Our MeDIP-seq analysis of genic region DNA methylation patterns revealed constitutively hypomethylated promoters of DEGs in SSC-enriched and progenitor-enriched spermatogonia (Figs. 2a,3k). We next mined our previously published datasets derived from regenerative SSC-enriched ID4-eGFP+/TSPAN8^High^ and progenitor-enriched ID4-eGFP+/TSPAN8^Low^ subpopulations of spermatogonia^28^, as well as published datasets derived from whole genome bisulfite sequencing analysis of subpopulations of “adult germline stem cell Thy1+” (AGSC Thy1) and “adult germline stem cell Kit+” (AGSC Kit) cells from adult testes^43, 44^ (Fig. 5). Overall, CpG sites were highly methylated genome-wide in all four spermatogonial subpopulations (approximately 90% CpGs methylated in each case) (Fig. 5a). 5’-regulatory regions showed wide variation in levels of DNA methylation that could be further subdivided into hypomethylated CpG-island-containing promoters and hypermethylated non-island promoters (Fig. 5a). Distal intergenic CpG islands were also hypomethylated. Differences in DNA methylation patterns between SSC-enriched and progenitor-enriched spermatogonial subpopulations were most evident within low methylated regions (LMRs) in which 10-50% of CpGs were methylated, or unmethylated regions (UMRs) in which <10% of CpGs were methylated^45^ (Figs. 5a-c, S5). Intergenic DNA methylation patterns in SSC-enriched ID4-eGFP+/TSPAN8^High^ and progenitor-enriched ID4-eGFP+/TSPAN8^Low^ subpopulations at P6 showed more similarity to one another than did SSC-enriched AGSC Thy1 and progenitor/differentiating spermatogonia-enriched AGSC Kit subpopulations in the adult testis (Fig. 5c-f). Promoter DNA methylation patterns showed predominant segregation into either hypermethylated or hypomethylated patterns, with very few showing intermediate level methylation (Fig. 5g-i). Spermatogonial subtype-specific LMRs aligned with enhancers, and, in many cases, correlated with subtype-specific distinctions in enhancer activity. Thus, the presence of an LMR typically coincided with the presence of one or more active or primed enhancers as well as with a prevalence of binding motifs for CTCF, CTCFL, and ELK1 (see below)^46^.

**Fig. 5:**
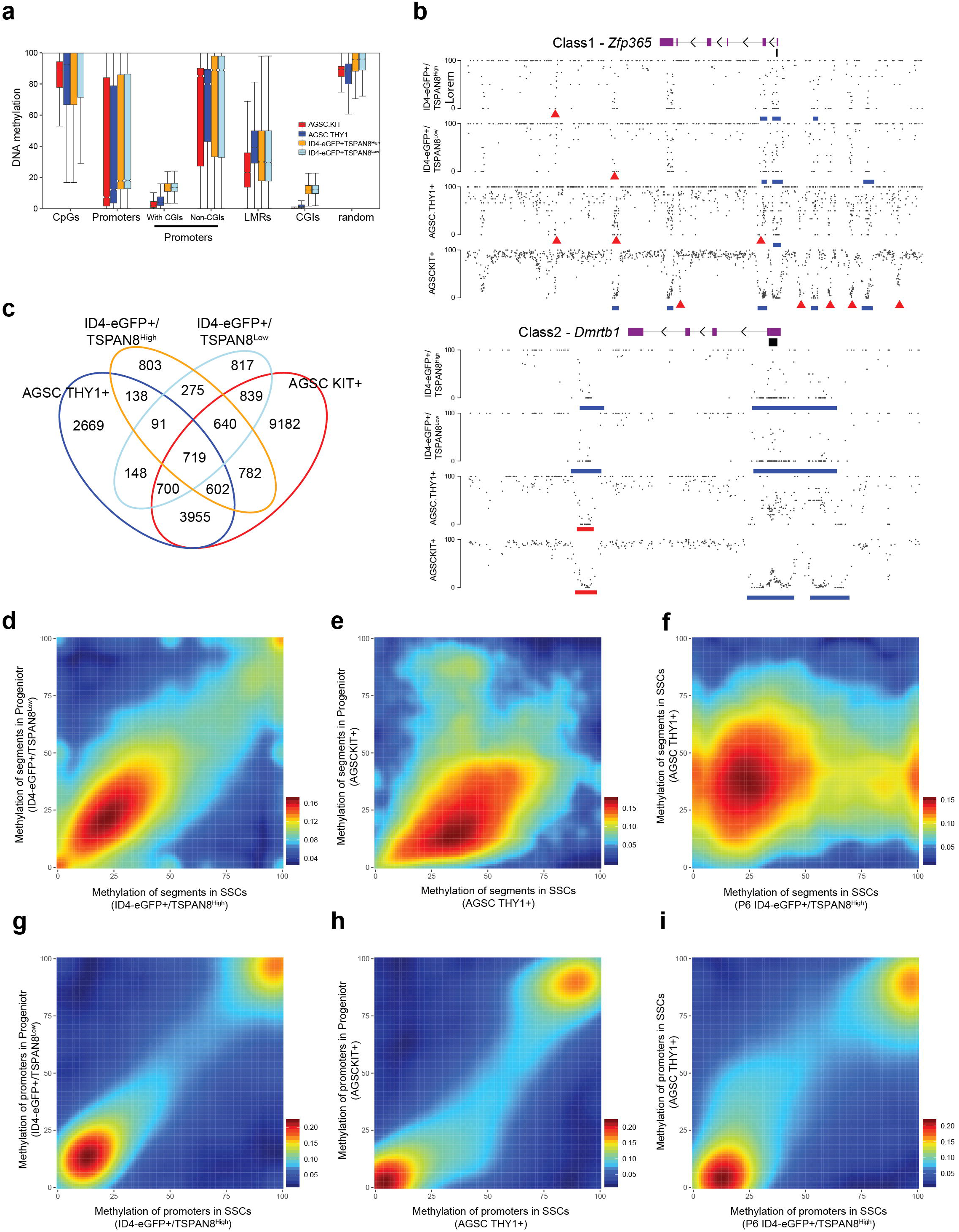
DNA methylation profiles in SSC-enriched ID4-eGFP+/TSPAN8^High^ and progenitor-enriched ID4-eGFP+/TSPAN8^Low^ spermatogonia from the immature (P6) mouse testis, and in SSC-enriched Thy1+ and progenitor/differentiating spermatogonia-enriched cKIT+ cells from the adult testis^43^. **a** DNA methylation levels associated with distinct genomic features including individual CpG dinucleotides (CpGs), gene promoters in general, CpG-island containing gene promoters, non-CpG island containing gene promoters, low methylated regions (LMRs), CpG islands (CGIs), and random regions of the genome in each spermatogonial subpopulation. **b** Exemplary genomic browser screenshots of DNA methylation levels in one Class 1 gene (*Zfp365*) and one Class 2 gene (*Dmrtb1*) in each of the four spermatogonial subpopulations. Each dot is a single CpG dinucleotide. Red triangles/bars demarcate LMRs, while blue bars indicate unmethylated regions (UMRs). **c** Venn diagram showing LMRs unique to each spermatogonial subtype (THY1+ or cKIT+ cells from the adult testis or ID4-eGFP+/TSPAN8^High^ or ID4-eGFP+/TSPAN8^Low^ spermatogonia from the immature [P6] testis), or common to each possible pair, trio or quartet of cellular subpopulations. **d-f** Smoothed scatter plots showing the extent of similarities or distinctions in patterns of DNA methylation in genomic segments in pairwise combinations of spermatogonial subpopulations including P6 ID4-eGFP+/TSPAN8^Low^ vs P6 ID4-eGFP+/TSPAN8^High^ (d), adult cKIT+ vs adult Thy1+ (e), adult Thy1+ vs P6 ID4-eGFP+/TSPAN8^High^ (f) cells. **g-i** Smoothed scatter plots showing the extent of similarities or distinctions in patterns of DNA methylation in gene promoters in pairwise combinations of spermatogonial subpopulations including P6 ID4-eGFP+/TSPAN8^Low^ vs P6 ID4-eGFP+/TSPAN8^High^ (g), adult cKIT+ vs adult Thy1+ (h), adult Thy1+ vs P6 ID4-eGFP+/TSPAN8^High^ (i) cells.

Interestingly, more differentially hypermethylated sites were found genome-wide in progenitor-enriched spermatogonia (Fig. S5a). Many of these were associated with enhancer functions such as stem cell maintenance, stem cell proliferation, cell-type specific development or differentiation, and homeostasis (Fig. S5b), consistent with down-regulation of SSC-specific genes. LMRs occurred primarily in distal intergenic regions (Fig. S5c), whereas UMRs occurred primarily in promoter regions (Fig. S5d). LMRs correlated with peaks of HeK4me1,2,3 and H3K27ac or H3K27me3, as well as with peaks of enhanced chromatin accessibility in both spermatogonial subtypes (Fig. S5e).

### Motif Enrichment Analysis Reveals Potential Regulators of Differential Epigenetic Programming Associated with Distinct Spermatogonial Subtypes

We performed motif enrichment analysis of promoter and enhancer regions associated with genes expressed differentially (Class 1,2) or constitutively (Class 3) in each spermatogonial subpopulation (Fig. 6). We found statistically significant (*p*<0.05) over-representation of binding motifs for 48 different factors in Class 1,2,3 gene promoters (Fig. 6a), a majority of which were enriched in promoters active in both spermatogonial subtypes. However, we observed differential enrichment of motifs for five factors (CDX4, HOXB4, EGR1, FOXA2, and ZIC1 [red triangles in Fig. 6a]) in promoters of Class 1 genes, and three factors (MAZ, ATOH1, and CTCFL [blue triangles in Fig. 6a]) in promoters of Class 2 genes. Coincidentally, transcripts encoding CDX4, HOXB4, FOXA2, and ZIC1 were up-regulated in SSC-enriched spermatogonia (as previously reported for *Zic1* ^21^), and transcripts encoding CTCFL were up-regulated in progenitor-enriched spermatogonia. Two of the factors for which we observed enriched binding motifs in Class 1 gene promoters, ZIC1 and EGR1, have been shown to promote self-renewal of SSCs^47^, and one for which we observed enriched motifs in Class 2 gene promoters (CTCFL) has been reported to promote spermatogonial proliferation and differentiation^48^.

**Fig. 6:**
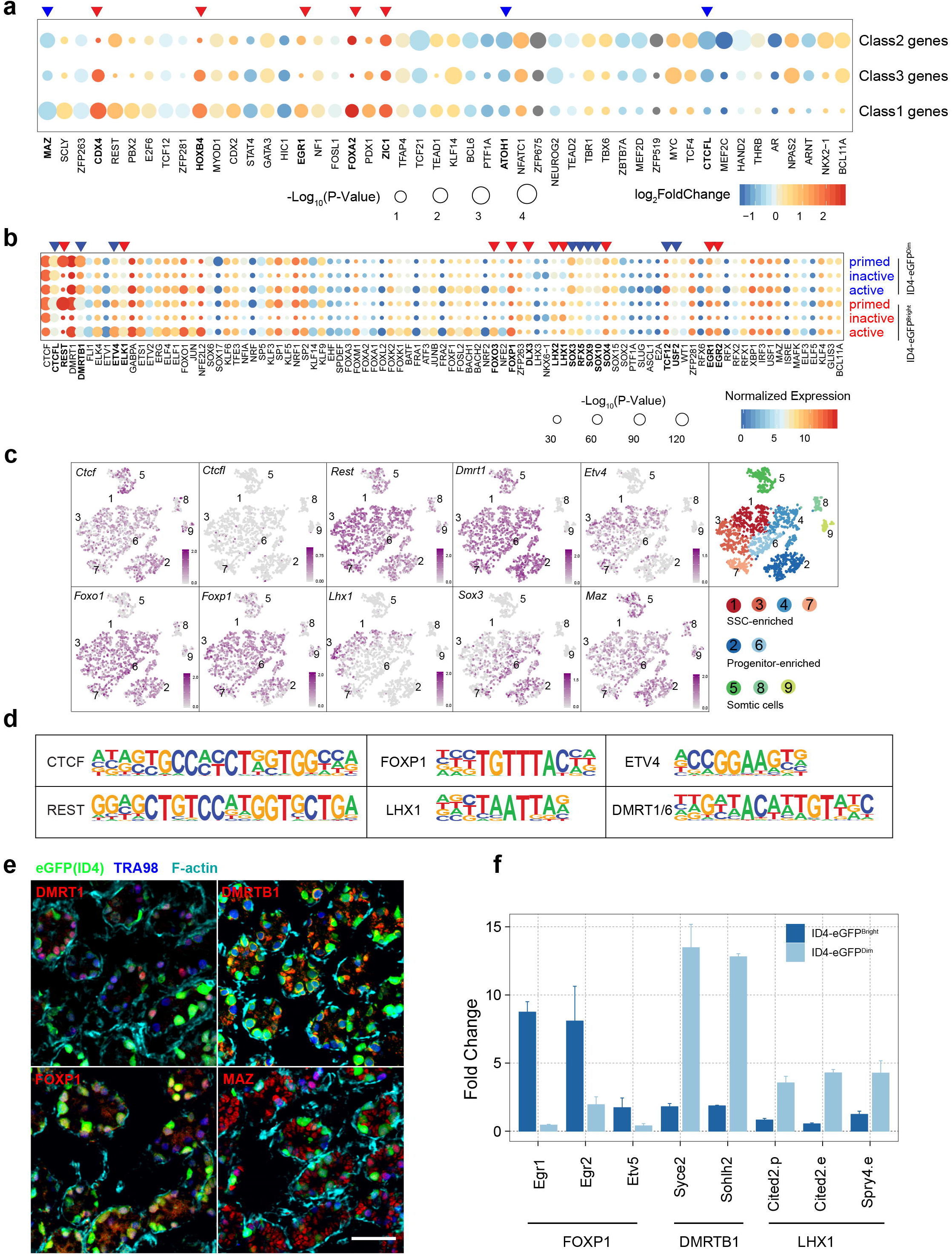
Predicted regulators of differential epigenetic programming in ID4-eGFP^Bright^ and ID4-eGFP^Dim^ spermatogonia. **a** Analysis of enrichment of transcription factor binding motifs within promoters of differentially and similarly expressed genes in SSC- and progenitor-enriched spermatogonial subpopulations. Each dot represents a corresponding motif, and the differential expression of the corresponding transcriptional regulator is shown as a color-keyed indication of Log Fold difference in enrichment of each motif in each gene promoter in each spermatogonial subtype (blue to red). The size of dots indicates the significance of each motif. **b** Analysis of enrichment of transcription factor binding motifs within primed, active or inactive enhancers associated with genes differentially expressed in SSC- and progenitor-enriched spermatogonial subpopulations. Each dot represents a motif, and the normalized expression of the corresponding transcriptional regulator is shown in different colors (green to red). The size of dots indicates the significance of each motif. **c** scRNA-seq data describing expression of mRNAs encoding exemplary transcription factors for which differential enrichment of binding motifs was detected in ID4-eGFP^Bright^ and ID4-eGFP^Dim^ spermatogonia, respectively. **d** Consensus sequences for exemplary factor-specific binding motifs. **e** Immunochemistry staining of DMRT1, DMRTB1, FOXP1 and MAZ in seminiferous cords in whole mount sections of P6 mouse testes. Green = expression of the ID4-eGFP^Bright^ or ID4-eGFP^Dim^ spermatogonial subtype marker transgene, dark blue = the germ cell marker, TRA98, and light blue = F-actin which delineates each seminiferous cord. **f** Gene- and factor-specific ChIP to examine differential binding of specific transcription factors to enhancers of differentially expressed genes in each spermatogonial subtype predicted by motif enrichment analysis. Each of three different factors shows significant differences in binding to the same motifs in ID4-eGFP^Bright^ and ID4-eGFP^Dim^ spermatogonia, with FOXP1 showing elevated binding to three target Class 1 genes in ID4-eGFP^Bright^ spermatogonia, and DMRTB1 and LHX1 showing elevated binding to two (DMRTB1) or three (LHX1) target Class 2 genes in ID4-eGFP^Dim^ spermatogonia.

Enhancers were characterized as regions of elevated chromatin accessibility containing hypomethylated DNA plus either H3K4me1+/H3K27ac+ (active enhancers), H3K4me1+/H3K27me3-/H3K27ac- (primed enhancers), or H3K4me1+/H3K27me3+ (inactive enhancers) histones (Fig. 6b). Enhancers were significantly more numerous and variably programmed than promoters of DEGs in ID4-eGFP^Bright^ and ID4-eGFP^Dim^ spermatogonia. We found significant enrichment (*p*<0.01) of binding motifs for >200 different factors within putative enhancer regions, of which 93 showed a -Log_10_(*p*-Value)>30 (Fig. 6b). These included binding motifs for several constitutive factors expressed at similar levels and enriched in enhancers in all states – active, primed or inactive – in both spermatogonial subpopulations. These were typically binding sites for factors involved in ubiquitous transcription complex formation and initiation of transcription, including SP1, SP2, MAZ, ELF1, ELF4, NRF1, KLF4 and IRF3 (Figs. 6b, S6).

Binding motifs for CTCF showed similar patterns in all types of enhancers in both spermatogonial subpopulations. Enrichment of binding sites for CTCFL, a testis-specific paralog of CTCF^49^, was highest in primed enhancers in both spermatogonial subpopulations, though levels of *Ctcfl* mRNA were higher in progenitor-enriched spermatogonia. The binding motif for REST was particularly enriched in primed enhancers, whereas that for DMRT1 was enriched in both active and primed enhancers in both spermatogonial subpopulations. Previously published DMRT1 ChIP-seq data from adult mouse testis showed that DMRT1 is bound to promoters of *Dusp6, Tlr3,* and *Ptpn9*^50^. *Dmrtb1* (aka *Dmrt6*) mRNA levels were higher in progenitor-enriched spermatogonia (Fig. 6b,c). Consensus enhancer binding motifs for six factors that were differentially enriched in ID4-eGFP^Bright^ and ID4-eGFP^Dim^ spermatogonia, CTCF, FOXP1, ETV4, REST, LHX1 and DMRT1/6, are shown in Figure 6d. Our IIF analysis indicated DMRT1 and DMRTB1 were more prevalent in progenitor than SSC nuclei (Fig. 6e). A previous study showed that DMRTB1 can repress expression of spermatogonial-expressed genes (*Dmrt1, Sohlh1, Sohlh2* and *Egr1*), and promote expression of meiotic genes (*Sycp2, Piwil2*)^51^. Our ChIP-qPCR data confirmed that DMRTB1 was differentially bound to enhancers of two Class 2 genes (*Sohlh2, Sycp2*) in progenitor-enriched spermatogonia where these genes were up-regulated (Fig. 6f).

Active enhancers were enriched for binding motifs for members of the FOX transcription factor family (FOXF1, FOXK1, FOXK2, FOXM1, FOXO1, FOXO3, FOXP1) in both spermatogonial subpopulations, and of these, transcripts encoding FOXO1 were most abundant (Fig. 6b). Expression of *Foxf1* is known to be restricted to germ cells in the testis (Fig. S6), and FOXF1 is thought to be a pioneer factor that directly regulates expression of *Etv1/4* and *Kit*^52^. FOXM1 is a transcriptional activator involved in regulating cell proliferation and maintenance of stem cells^53, 54^. The FOXM1 binding motif was more enriched in active enhancers in SSC-enriched spermatogonia (Fig. 6b). FOXO1 and FOXO3 have been reported to regulate *Ret* to maintain SSC self-renewal^55^, and binding motifs for these factors were also more enriched in active than in primed or inactive enhancers in both spermatogonial subpopulations. FOXP1 binding motifs were also found to be prevalent at active enhancers in both spermatogonial subpopulations, and expression of *Foxp1* was elevated in spermatogonia in general (Fig. 6c). FOXP1 binding motifs have also been shown to be enriched in human SSCs^56^, and FOXP1 has been implicated as a regulator of cell fate in multiple other types of stem cells^57, 58, 59, 60^. Our IHC data showed FOXP1 is selectively localized in nuclei of SSC-enriched ID4-eGFP^Bright^ spermatogonia (Fig. 6e), and our ChIP-qPCR data showed differential binding of FOXP1 to active enhancers of three Class 1 target genes, *Egr1, Egr2,* and *Etv5*, specifically in SSC-enriched spermatogonia (Fig. 6f). By contrast, MAZ was prevalent in somatic Sertoli cells as well as in progenitor spermatogonia, but appeared to be excluded from nuclei of ID4-eGFP^Bright^ spermatogonia (Fig. 6e).

Members of the SOX family (SOX3, SOX4, SOX10 and SOX15), which are known to promote cellular differentiation and cell fate determination ^61^, showed elevated binding motif enrichment in primed and active enhancers in progenitor-enriched spermatogonia (Fig. 6b). SOX3 has been reported to colocalize with NEUROG3 and is specifically expressed in proliferating spermatogonia^62^, though enrichment of binding motifs for SOX3 has also been reported in human SSCs^56^. Both our bulk RNA-seq (Fig. S6) and previous scRNA-seq data^21^ showed elevated *Sox3* transcripts in progenitor-enriched spermatogonia.

We observed higher enrichment of binding motifs for LHX1, LHX2, LHX3 and DLX3 in inactive enhancers in both SSC-enriched and progenitor-enriched spermatogonia. Interestingly, levels of *Lhx1* mRNA were higher in SSC-enriched spermatogonia than in progenitor-enriched spermatogonia, but LHX1 was more robustly bound to enhancers of certain down-regulated Class 1 genes in progenitor-enriched spermatogonia (Fig. 6f). Thus, binding of LHX1 may repress expression of Class 1 genes such as *Cited2* and *Spry4*. Finally, as expected, we detected no or extremely low expression of many factors known to specifically regulate differentiation of various somatic cell types, including ATOH1, CDX2, CDX4, ELF5, ISRE, RFX, RFX6, E2A, SLUG, LHX3, NRF, NRF2, FRA1, FRA2, FOXL2, EHF, SOX17, STAT4, HAND2, ZFP519, NEUROG2, ZFP675, and NKX2-1.

### Differential Fates of ID4-eGFP^Bright^ and ID4-eGFP^Dim^ Spermatogonia Are Associated with Coordinated, Multiparametric Programming of Differentially Expressed Genes

Ultimately, it is specific <u>combinations</u> of chromatin states defined by epigenetic signatures and specific transcription factor interactions that drive differential gene expression, which, in turn, establishes distinct fates of different cell types or subtypes^63^. Thus, we integrated our ChIP-seq, MeDIP-seq, ATAC-seq and RNA-seq data using multivariate Hidden Markov Model building by ChromHMM to identify and characterize 15 different chromatin states within the spermatogonial genome in agreement with ENCODE project methods (Fig. 7a). These 15 different states were assessed in SSC- and progenitor-enriched spermatogonia to compare the extent to which transitions among each were associated with unique fates of spermatogonial subtypes (Fig. 7b). For instance, the pattern of quiescent enhancers in state 7 showed little or no difference between ID4-eGFP^Bright^ and ID4-eGFP^Dim^ spermatogonia, likely representing enhancers involved with gene expression in non-spermatogonial cell types. In contrast, a portion of enhancers displaying either an inactive (state 1) or active (state 6) status in SSC-enriched spermatogonia resolved to a quiescent state (state 5) in progenitor-enriched spermatogonia (Fig. 7b). Interestingly, in both ID4-eGFP^Bright^ and ID4-eGFP^Dim^ spermatogonia, bivalent enhancers (state 10) were closely associated with repressed (state 9) or poised (state 11) promoters.

**Fig. 7:**
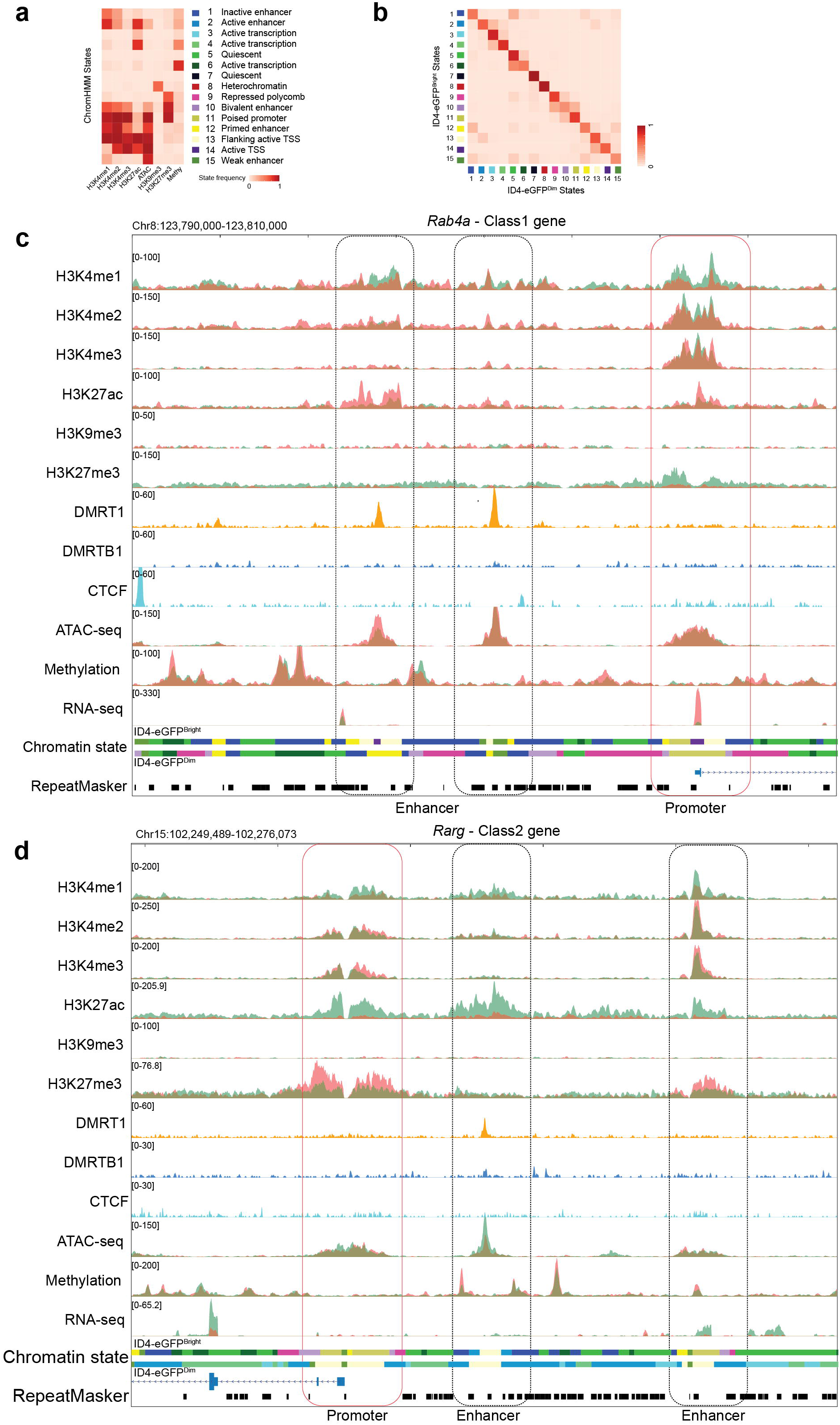
Coordinated epigenetic programming regulates differential gene expression in ID4-eGFP^Bright^ and ID4-eGFP^Dim^ spermatogonia. **a** 15 distinct chromatin states predicted by ChromHMM analysis of eight different epigenetic parameters. **b** Spermatogonial-subtype specific transitions among chromatin states. **c,d** Genome browser views of an exemplary Class 1 gene (*Rab4a*) (c) and Class 2 gene (*Rarg*) (d) showing relative abundance of: transcripts detected by bulk RNA-seq; each of eight different epigenetic parameters (H3K4me1-3, H3K9me3, H3K27ac, & H3K27me3 detected by ChIP-seq; chromatin accessibility detected by ATAC-seq; and DNA methylation detected by MeDIP-seq) in ID4-eGFP^Bright^ and/or ID4-eGFP^Dim^ spermatogonia from the P6 testis; plus peaks indicative of binding of DMRT1, DMRTB1 and CTCF in adult testis cells mined from published datasets, and the chromatin states each combination of these parameters indicates in each region of each gene, color-coded to match the states defined in (a). Bulk RNA-seq, histone modification-specific ChIP-seq, ATAC-seq and MeDIP-seq analyses were each carried out on purified populations of ID4-eGFP^Bright^ and ID4-eGFP^Dim^ spermatogonial subtypes and the data from each subtype is shown as overlays with data from ID4-eGFP^Bright^ spermatogonia shown as coral colored tracings and that from ID4-eGFP^Dim^ spermatogonia shown as light green tracings, and that from overlapping tracings from both spermatogonial subtypes shown as a dark olive-brown color. Each browser view shows the entire sequence of each gene including promoter(s) and enhancers as noted.

Parallel visualization of tracks indicative of read intensities derived from each epigenomic analysis facilitated the most direct, comprehensive, locus-specific comparisons of coordinated epigenetic programming in SSC- and progenitor-enriched spermatogonia, respectively (Figs. 7c,d). This revealed unique epigenetic signatures associated with promoters or enhancers of DEGs in these two spermatogonial subtypes. Thus, as noted above, programming patterns associated with promoters of DEGs up-regulated in one or the other spermatogonial subpopulation included enriched H3K4me1,2,3 + H3K27ac, depleted H3K27me3, hypomethylated DNA and elevated chromatin accessibility (Figs. 7c,d). Interestingly, promoters of Class 1 or 2 genes showed enriched H3K4me1,2,3, hypomethylated DNA and elevated chromatin accessibility in both spermatogonial subpopulations, despite the fact that these genes were differentially expressed in each. However, enrichment of H3K27ac or H3K27me3 varied directly with up- or down-regulation of genes in each subpopulation, respectively. This suggests differential enrichment of these two promoter region modifications contributes directly to differential regulation of Class 1 and 2 genes in SSCs versus progenitors. Enhancers of DEGs showed enrichment of H3K27ac + depletion of H3K4me1 for up-regulated genes and enrichment of H3K27me3 + H3K4me1 for down-regulated genes (Figs. 7c,d).

We augmented these data with those from published reports of genome-wide binding patterns of DMRT1^64^, DMRTB1^51^, and CTCF^65^ in adult testis tissue. Although no distinction was made between SSC- and progenitor-enriched spermatogonia, or even between spermatogenic and somatic cells in these studies, it is noteworthy that peaks of DMRT1 and DMRTB1 binding were detected at enhancers of many of the Class 1 and Class 2 genes identified in our study (Figs. 7c,d). Binding motifs for CTCF were observed in distal flanking intergenic regions consistent with reports that CTCF plays an important role in 3-dimensional organization of the genome to mediate long range enhancer-promoter interactions contributing to cell fate determination^66^. Finally, using the code shown in Figure 7a, we were able to predict the arrangement of chromatin states in a linear context in each spermatogonial subtype (Figs. 7c,d), as well as the extent to which these states varied either between Class 1 and Class 2 genes within each spermatogonial subtype, or within Class 1 or Class 2 genes between spermatogonial subtypes (Figs. 7c,d). Additional data regarding the genome-wide distribution of potential regulatory elements and chromatin states identified by our ChromHMM analysis are shown in Figure S7.

## Discussion

The ENCODE^67^, NIH Roadmap Epigenomics^68^ and related^35^ studies characterized the presence and variable states of key regulatory elements throughout the genomes of >100 different somatic cell types on the basis of multiparametric integrative analysis methodology ^63^, but did not examine any germ cell types. One previous study provided an initial characterization of histone modifications in fetal mouse germ cell types, but did not examine postnatal germ cells, and therefore did not characterize epigenetic programming distinguishing SSCs from progenitors. Fetal and postnatal spermatogenic cell types have also been assessed for poised genes but those studies were limited to a limited set of histone modifications^69^. Results from transplantation studies have shown that regenerative SSC capacity resides in only a small subpopulation of undifferentiated spermatogonia^20^, however the advent of the *Id4-eGfp* transgenic mouse has facilitated selective recovery of spermatogonial subpopulations highly enriched for, or significantly depleted of this capacity^20, 27^. Multiple recent studies established consistent differences in gene expression patterns in SSC- and progenitor-enriched subpopulations^9, 20, 21, 22, 23, 28^, suggesting these spermatogonial subtypes represent the emergence of distinct cell fates driven by distinct transcriptomes. Here, we have extended these analyses by conducting the first comprehensive, multiparametric integrative epigenomic analysis of epigenetic programming associated with spermatogonial-subtype specific DEGs in a manner similar to that previously reported for somatic cell types [78,79,37]. This revealed distinct epigenetic landscapes specifically associated with differential gene expression in the two spermatogonial subtypes, which we then further mined to identify binding sites for specific factors that may either direct establishment of this differential epigenetic programming or mediate its effects to coordinate subsequent differential expression of genes required to either maintain SSC fate or initiate progenitor fate.

Given the common developmental ancestry of SSCs and progenitors, it is not surprising that they display predominantly similar transcriptomes as evidenced by equivalent transcript levels for 95% of genes expressed in either ID4-eGFP^Bright^ or ID4-eGFP^Dim^ spermatogonia, or both. However, we did detect differential expression of a substantial number of genes distinguishing the two spermatogonial subpopulations on the basis of scRNA-seq and bulk RNA-seq, with 669 genes up-regulated in ID4-eGFP^Bright^ cells and 373 genes up-regulated in ID4-eGFP^Dim^ cells, consistent with previously reported results^9, 20, 21, 28^.

Chromatin states are an inherent, biologically-informative feature of the genome that are often cell-type or -subtype specific^35^. The majority of epigenetically dynamic regions identified throughout the genomes of many different somatic cell types have been found in distal intergenic regions, consistent with the notion that differential expression of protein-encoding genes is regulated most precisely by intergenic enhancers^32, 41, 67^. Our results indicate this observation can now be extended to DEGs in spermatogonial subtypes as well. Thus, genes expressed at similar levels in the two spermatogonial subpopulations showed little or no detectable differences in epigenetic programming, while DEGs showed specific distinctions in certain epigenetic parameters. In particular, differential enrichment of H3K27ac or H3K27me3 in promoter regions, and of H3K27ac, H3K27me3 and/or H3K4me1 at enhancers, correlated with up- or down-regulated transcript levels in each spermatogonial subtype. We also found that differentially programmed enhancers typically overlapped with differences in patterns of partially methylated regions in distal intergenic areas. Thus, spermatogonial subtype-specific gene expression patterns appear to be regulated by differential patterns of specific histone modifications and DNA methylation at intergenic enhancers. However, several other chromatin parameters, including patterns of H3K4me1,2,3, chromatin accessibility and DNA methylation at promoters, and those of H3K4me2,3 and chromatin accessibility at enhancers, showed no significant variation among DEGs regardless of whether the gene was expressed in both or only one spermatogonial subpopulation. Collectively, this is consistent with the notion that development of distinct cell fates from a similar precursor cell type involves an ordered series of changes in epigenetic programming to first initiate and subsequently stabilize differential gene expression associated with distinct cell types.

Previous reports have described epigenetic poising of genes (promoters simultaneously marked with H3K4me3 and H3K27me3) in the spermatogenic lineage that appears to predispose the capacity of the paternal genome to rapidly transition to an embryonic transcriptome following fertilization^69^. We found that many non-poised genes expressed in SSC-enriched spermatogonia become poised and repressed in progenitor-enriched spermatogonia. This raises the intriguing possibility that, in addition to marking initiation of commitment to the spermatogenic differentiation pathway, the SSC-progenitor transition also demarcates initiation of a final phase of epigenetic programming to prepare the paternal genome for post-fertilization functions.

Enhancers and partially methylated regions are both rich in transcription factor binding sites^45^, and it has been shown that transcription factors act as key drivers of differential states of activity or inactivity at enhancers^70^. Our motif enrichment analysis revealed many binding sites common to regulatory regions in both spermatogonial subtypes. However, we also identified differential enrichment of certain motifs in promoters or enhancers regulating DEGs in ID4-eGFP^bright^ and ID4-eGFP^dim^ spermatogonia, and these formed the basis for testable predictions of differential binding of specific transcription factors in each spermatogonial subpopulation. We confirmed these predictions for three such factors – FOXP1, DMRTB1 and LHX1 – each of which showed spermatogonial subpopulation-specific differences in a) expression at the RNA level, b) prevalence/intracellular location at the protein level, and/or c) binding to enhancers of differentially expressed target genes.

Our results do not unequivocally resolve the differing theories regarding the developmental dynamics affecting SSCs and progenitors – particularly the question of whether or not these are equipotent spermatogonial subtypes that continually interconvert during steady-state spermatogenesis as some have suggested^17, 71^ and others have questioned^2^. A combination of further assessments of spermatogonial subpopulations showing directly testable enhanced or depleted representation of regenerative SSCs based on the spermatogonial transplantation assay, along with appropriate lineage-tracing and ablation studies will be required to reach a definitive resolution of this question. However, we have identified distinct epigenetic programming characteristics associated with differential gene expression patterns distinguishing regenerative SSC-rich and non-regenerative progenitor-rich spermatogonial subpopulations. We suggest this differential epigenetic programming drives the initiation of cell fate divergence between SSCs and progenitors, thereby directing a significant developmental switch between retention of SSC fate and initiation of spermatogenic differentiation, respectively.

Finally, we previously suggested that the initial, foundational pool of SSCs that forms in the postnatal mouse testis may derive from a distinct subpopulation of prospermatogonia that become uniquely programmed during late fetal and early postnatal stages such that they are predetermined to form the foundational SSCs^8, 9, 21, 26, 28, 72^. Our characterization of the epigenetic landscape within foundational SSCs now provides the first insight into the type of epigenetic programming that may underpin such a predetermination mechanism.

## Methods

### Mice and cells

All experiments utilizing animals were preapproved by the Institutional Animal Care and Use Committee of the University of Texas at San Antonio (Assurance A3592-01) and were performed in accordance with the National Institutes of Health (NIH) *Guide for the Care and Use of Laboratory Animals*. Testes were recovered from 6-day old (P6) F1 male offspring of a cross between *Id4-eGfp* (LT-11B6) and either C57Bl6/JJ or *Rosa26-lacZ* (The Jackson Laboratory #000664, #002073) mice and used to generate suspensions of cells by enzymatic digestion as described ^28^. Briefly, ID4-eGFP+ testes were distinguished by fluorescence microscopy and then subjected to dissociation and FACS sorting. After removing the tunica albuginea, testes were digested with DNAaseI + trypsin (0.05% trypsin, 10 µg/ml Dnase I in HBSS with 0.5 mM EDTA) to generate a single cell suspension. The resulting dissociated cells were washed and resuspended in Dulbecco’s phosphate-buffered saline (**DPBS**) + 10% FBS, filtered through a 40 micron strainer to remove Sertoli cells and cell clumps prior to being subjected to fluorescence-activated cell sorting (**FACS**) as previously described^28^. Cells at a concentration of approximately 15×10^6^ cells/ml DPBS + 10% FBS were subjected to flow cytometry using BD FACS Aria. Propidium iodide (**PI**) was added to discriminate dead cells at 5µl/10^6^ cells. Positive ID4-eGFP epifluorescence was determined by comparison to testis cells from testes of wild-type mice lacking the P6 *Id4-eGfp* transgene. The gating area of eGFP positive was subdivided into thirds to define the ID4-eGFP+ subsets as being Dim (lower third) or Bright (upper third) by fluorescent intensity as described^20, 21^.

### Bulk RNA-seq

Aliquots of cells from each of four replicate samples of each spermtogonial subpopulation were used for separate bulk RNA-seq analyses to catalogue gene expression in each spermatogonial subtype. Populations of at least ≥1000 ID4-eGFP^Bright^ or ID4-eGFP^Dim^ cells recovered by FACS sorting were counted, pelleted, and subjected to direct cDNA synthesis using the SMART-Seq v4 ultra Low Input RNA Kit for Sequencing (Clontech Laboratories #634888). Approximately 250pg of cDNA was used for preparation of dual-indexed libraries using the Nextera XT DNA Library Preparation Kit (Illumina # FC-131-1002) following the manufacturer’s procedures.

### ChIP-seq

Aliquots of cells from each of four replicate samples of each spermtogonial subpopulation were used for separate ChIP-seq analyses to detect genome-wide enrichment patterns of six different histone modifications – H3K4me1, H3K4me2, H3K4me3, H3K9me3, H3K27me3 and H3K27ac. Approximately 1×10^6^ cells were used for each immunoprecipitation (**IP**). ULI-NChIP-seq was performed as previously described^73^. Briefly, FACS-sorted cells were pelleted and re-suspended in nuclear isolation buffer (Sigma #NUC101-1KT). Depending on input size, chromatin was fragmented for 5-7.5 min using MNase, and diluted in NChIP immunoprecipitation buffer (20mM Tris-HCl pH 8.0, 2mM EDTA, 15 mM NaCl, 0.1% Triton X-100, 1×EDTA-free protease inhibitor cocktail and 1mM phenylmethanesulfonyl fluoride). 10% of each sample was reserved as input control. Chromatin was pre-cleared with 5 or 10 µl of 1:1 protein A:G Dynabeads (Life Technologies #10015D) and immunoprecipitated with H3K4me1 (Abcam #ab8895), H3K4me2 (Abcam #ab7766), H3K4me3 (Abcam #ab8580), H3K9me3 (Abcam #ab8898), H3K27ac (Active Motif #39133) and H3K27me3 (Abcam #ab6002) antibody-bead complexes overnight at 4°C. IPed complexes were washed twice with 400 µl of ChIP wash buffer I (20 mM Tris-HCl, pH 8.0, 0.1% SDS, 1% Triton X-100, 0.1% deoxycholate, 2 mM EDTA and 150mM NaCl) and twice with 400 µl of ChIP wash buffer II (20mM Tris-HCl (pH 8.0), 0.1% SDS, 1% Triton X-100, 0.1% deoxycholate, 2mM EDTA and 500mM NaCl). Protein-DNA complexes were eluted in 30 µl of ChIP elution buffer (100mM NaHCO_3_ and 1% SDS) for 2h at 68 °C. IPed material was purified by PCI, ethanol-precipitated and raw ChIP material was resuspended in 10 mM Tris-HCl pH 8.0. Fragment length was confirmed using a Bioanalyzer (Aglient Technology), and DNA concentration was determined by the Qubit dsDNA HS Assay Kit (Invitrogen #Q32854). Illumina libraries were constructed using a modified custom paired-end protocol. In brief, samples were end-repaired in 1× T4 DNA ligase buffer, 0.4 mM dNTP mix, 2.25U T4 DNA polymerase, 0.75U Klenow DNA polymerase and 7.5U T4 polynucleotide kinase for 30 min at 21-25°C, then A-tailed in 1× NEB buffer 2, 0.4 mM dNTPs and 3.75U of Klenow(exo-) for 30 min at 37°C and then ligated in 1× rapid DNA ligation buffer plus 1mM Illumina PE adapters and 1,600U DNA ligase for 1-8h at 21-25°C. Ligated fragments were amplified using dual-indexed primers for Illumina (NEB #E7600S) for 8-10 PCR cycles. DNA was purified with 1.8× volume Ampure XP DNA purification beads (Beckman Coulter #A63881) between each step. Fragment length was again checked by Bioanalyzer (Aglient Technology), and DNA concentration was determined using the Qubit dsDNA HS Assay Kit (Invitrogen #Q32854).

### ATAC-seq

After FACS sorting, each aliqout of fresh cells (∼50,000 cells/aliquot) was pelleted and re-suspended in transposition mix (25µl 2x TD buffer, 2.5 µl Tn5 transposase (100 nM final), 16.5µl PBS, 0.5 µl 1% Digitonin, 0.5µl 10% Tween-20, 5µl H_2_O) and incubated at 37°C for 30 min in a thermomixer. The mix was then treated with a Zymo DNA Clean and Concentrator-5 Kit (Zymo research #D4014). ATAC-seq libraries were constructed in the same manner as that described above for ChIP-seq libraries.

### MeDIP-seq

MeDIP-seq libraries were constructed as previously described^74^. After FACS sorting, each aliqout (∼50,000) of fresh cells was pelleted and re-suspended in lysis buffer (50mM Tris pH 8.0, 10 mM EDTA, 100 mM NaCl, 1% SDS, 0.5 mg/ml proteinase K) and incubated at 55°C for 5h. Genomic DNA was isolated using Phenol:Chloroform:Isoamyl Alcohol (Invitrogen #15593031), and sheared using a Bioruptor (Diagenode UCD-200). 10% raw sheared DNA was retained to serve as input control. Samples were end-repaired in 1x T4 DNA ligase buffer, 0.4mM dNTP mix, 2.25U T4 DNA polymerase, 0.75U Klenow DNA polymerase and 7.5U T4 polynucleotide kinase for 30 min at 21-25°C, then A-tailed in 1x NEB buffer 2, 0.4mM dNTPs and 3.75U of Klenow(exo-) for 30 min at 37°C, and then ligated in 1x rapid DNA ligation buffer, 1mM Illumina PE adapters and 1,600U DNA ligase for 1-8 h at 21-25°C. Samples were denatured at 95°C for 10 min, then transfered immediately to ice to prevent re-annealing. 0.2pM λ fragments (50% methylated) were used as a spike-in control. MeDIP on purified adapter-ligated DNA with spike-in was performed in 0.1 M Na_2_HPO_4_/NaH_2_PO_4_, 35μl of 2 M NaCl, 2.5 μ 10% Triton X-100 and 1μl of anti-methylcytidine antibody (1 mg/ml Diagenode # MAb-081-μ 100) overnight. DNA-IgG complexes were captured by protein A/G agarose beads. DNA was extracted by phenol:chloroform:isoamyl alcohol (25:24:1, v/v). Recovery (%) of MeDIP was calculated as 2^amplification efficiency^(Adjusted InputCt - MeDIPCt)^ × 100%. Specificity of MeDIP is calculated as: Specificity = 1-(unmeth recovery/meth recovery). Only libraries with specificity ≥95% and unmethylated recovery of < 1% were used for further analysis.

### Next Generation DNA Sequencing

Libraries were quantified by PCR using the NEBNext Library Quant Kit from Illumina (NEB #E7630L). After quantification, libraries were pooled in equal molar concentrations. RNA-seq libraries were sequenced on an Illumina HiSeq 2000 (PE100) at the University of Texas Southwestern Medical Center Sequencing Core. ChIP-seq, ATAC-seq and MeDIP-seq libraries were sequenced on an Illumina Hiseq 3000 sequencer (PE100) at the University of Texas Health Science Center at San Antonio Sequencing Core according to standard Illumina protocols.

### Bioinformatics Analyses

#### Sequencing and Alignments

All raw fastq files were mapped to the UCSC mm10 genome reference using Rsubread or QuasR^75, 76^

#### RNA-seq analysis

Count matrices assigned to genes were obtained using featureCounts^77^. Differential expression was inferred using DESeq2^78^. Genes with *p* < 0.01 and LFC >1.5 were considered significantly differentially expressed.

#### ChIP-seq analysis

Sites of differential histone modification were determined by a sliding window model and visualized by volcano plots, and sites displaying LFC >1.5, plus *p* < 0.01, and FDR < 0.01 were considered significantly differentially modified^79^. RPKM of histone H3 modifications including H3K4me1, H3K4me2, H3K4me3, H3K9me3, H3K27ac and H3K27me3 on promoters (TSS ± 500bp) were determined, log-transformed and defined as positive if their enrichment value was ≥ a threshold established by fitting a two-component Gaussian mixture model using Mclust^80^. Read coverage, K-means clustering and heatmap visualization were performed by deepTools and ngs.plot.r ^81, 82^.

#### ATAC-seq analysis

Differentially accessible chromatin sites were determined by a sliding window model and visualized by volcano plots, and those displaying log fold change >1.5, plus *p* value <0.01, and FDR <0.01 were considered as significantly differentially accessible^79^. To identify potential enhancer loci, sequence within +/- 1kb from each ATAC-seq peak was examined. All ATAC-peaks not overlapping with promoters, known gene bodies, or extended transcription end sites were examined. The histone enrichment in these regions was determined by fitting a two-component Gaussian mixture model using Mclust^80^.

#### MeDIP-seq analysis

Genome-wide differential coverage analysis of MeDIP-seq data was conducted using MEDIPS^83^. Differentially methylated regions were annotated by ChIPseeker and interpreted by GREAT.

#### Peak calling

Duplicated reads were removed by Picard (http://broadinstitute.github.io/picard/). Regions enriched for H3K9me3 or H3K27me3 were determined using MACS2 peak callers on non-duplicated, uniquely aligned reads. Broad peaks (H3K9me3, H3K37me3) were identified using MACS2 broadpeaks (*p* < 1×10^-6^, FDR<0.01) and narrow peaks (H3K4me1, H3K4me2, H3K4me3, H3K27ac, ATAC-seq and MeDIP-seq) were identified with MACS2 (*p* < 1×10^-6^, FDR< 0.01). Peaks closer than 2 kb apart were merged and peaks larger than 0.5 kb were included in our analysis^84^. Peaks were compared and annotated using ChIPseeker^85^.

#### Gene Ontology analysis

GSEA were determined using clusterProfiler^86^. GO analysis were determined using DAVID or clusterProfiler. Functional gene network analysis was conducted using FGNet^39^. Functional interpretation of enhancer-like regions was performed using GREAT using default parameters^87^.

#### Motif analysis

Enrichment analysis of known motifs within promoter and enhancer regions was analyzed with HOMER with default parameters and a fragment size of 200 bp. All known motifs used in our study were defined by HOMER.

#### Integrating chromatin states

Chromatin states, were assigned after the mouse genome was discretized into 200bp bins and subjected to a 15-state Hidden Markov modeling analyses using the ChromHMM method with default parameters^63^. CTCF^88^, DMRT1^64^ and DMRTB1^51^ ChIP-seq coverage from published studies of adult mouse testes and data from our analyses of P6 mouse testes were integrated and visualized by pyGenomeTracks and UCSC genome browser^89^.

### Factor/Gene-Specific ChIP and Real-time PCR

FACS-sorted populations of P6 ID4-eGFP^Bright^ and ID4-eGFP^Dim^ spermatogonia were fixed in freshly prepared cross-linking buffer (0.1M NaCl, 1mM EDTA, 0.5mM EGTA, 50mM HEPES (pH 8.0), 11% Formaldehyde). Cells were lysed in buffer L1 (140mM NaCl, 1mM EDTA, 50mM HEPES, 10% Glycerol, 0.5% NP-40, 0.25% Triton X-100) and nuclei were isolated using buffer L2 (200mM NaCl, 1mM EDTA, 0.5mM EGTA, 10mM Tris). Chromatin was sheared to average size of 500bp using a Bioruptor (Diagenode UCD-200). FOXP1 (Abcam # ab16645), DMRTB1 (Abcam # ab241275), LHX1 (Santa Cruz Biotechnology #sc-515631) and IgG (Abcam # ab37355 & ab171870) antibodies were coupled to DynBeads in DPBS (5 mg/ml BSA) by incubating overnight on a rotating platform at 4°C. Chromatin was precipitated by antibody-bead complexes in IP buffer (1% TritonX-100, 0.1% deoxycholate sodium salt, 1× Complete protease inhibitor, 10mM Tris-Cl (pH 8.0), 1mM EDTA) overnight on a rotating platform at 4°C. DNA-antibody-bead complexes were washed 10 times using freshly prepared RIPA buffer (50mM HEPES, 1 mM EDTA, 1% NP-40, 0.7% deoxycholate sodium salt, 0.5M LiCl, 1× complete protease inhibitor). DNA was eluted in elution buffer (10 mM Tris (pH 8.0), 1mM EDTA, 1% SDS) and cross-linkages were reversed overnight at 65°C. After proteinase K digestion, DNA was purified with PCI. ChIP-qPCR was performed on QuantStudio 3 instrument (Applied Biosystems) using Luna® Universal qPCR Master Mix (NEB #M3003S) following instructions in the reagent manual. ChIP DNA and control DNA were used as templates. Primers flanking potential factor-binding sites were designed by Primer-BLAST^90^. Fold Enrichment was calculated by 2^-ΔΔCt^.

### Indirect Immunofluorescence Microscopy

Immunolabeling was done as previously described^91^. Briefly, testes were immersion-fixed in fresh 4% paraformaldehyde, washed in PBS, incubated overnight in 30% sucrose at 4°C, and frozen in O.C.T. Five micrometer sections were incubated in blocking reagent (PBS containing 3% BSA and 0.1% Triton X-100) for 30 min at room temperature. Primary antibodies were used against DMRT1 (Abcam #ab222895), DMRTB1 (Abcam #ab241275), FOXP1 (Abcam #ab16645), or MAZ (Abcam #ab85725; all at 1:500). Primary antibodies were diluted with blocking reagent and incubated on tissue sections for 1 h at room temperature. Primary antibody was omitted as a negative control. Following stringency washes, sections were incubated in secondary antibody (1:500, AlexaFluor donkey anti-rabbit-555, Invitrogen) with phalloidin-405 (at 1:500, Invitrogen) for 1h at room temperature. Blocking and antibody incubations were done in PBS containing 3% BSA and 0.1% TritonX-100, and stringency washes were done with PBS and 0.1% TritonX-100. Cover slips were mounted with Vectastain containing DAPI (Vector Laboratories), and images obtained using a Fluoview FV1000 confocal laser-scanning microscope (Olympus America).

### Data availability

All datasets generated from this study have been archived in the NCBI GEO database, with the accession number GSE131657. P6 ID4-eGFP+ single cell RNA-seq data was obatian from GSE109049^21^. P6 spermatogonia RRBS data was obtained from GSE83311 and GSE83422^28^. The adult male germline stem cell BiSeq data was obtained from GSE49624^43^. Adult mouse testis ChIP-seq data for CTCF, DMRT1, and DMRTB1 was obtained from GSM918711^88^, GSE64892^64^, and GSM1480189^51^, respectively.

## Acknowledgements

The authors thank Mr. Sean Vargas for assistance with procedures conducted in the UTSA Genomics Core.

## Authors’ contributions

KC and JRM conceived and designed the experiments. KC performed all RNA-seq, ChIP-seq, ATAC-seq and MeDIP-seq experiments and analyzed all of the data from each of those experiments and prepared all of the figures except Fig. 6d. KC also performed the gene-specific ChIP experiments and the motif enrichment analyses. I-CC assisted with cell sorting. BJH and CG conducted indirect immunofluorescence microscopy and prepared Fig. 6d. JMO provided the Id4-egfp transgenic mouse line and related spermatogonial transplantation results. KC mined single-cell transcriptome data from datasets previously prepared by KC, BPH and JRM. BPH assisted with interpretations of the scRNA-seq data and directs the UTSA Genomics Core in which KC prepared libraries for sequencing. All authors contributed to interpretation of the data. JRM wrote the manuscript with very significant input from KC plus additional input from all other authors. All authors read and accepted the final version of the manuscript.

## References

1. Johnson L, Petty CS, Neaves WB. A Comparative-Study of Daily Sperm Production and Testicular Composition in Humans and Rats. Biology of Reproduction 22, 1233–1243 (1980).

2. de Rooij DG. The nature and dynamics of spermatogonial stem cells. Development 144, 3022–3030 (2017).

3. Brinster RL, Avarbock MR. Germline transmission of donor haplotype following spermatogonial transplantation. Proc Natl Acad Sci U S A 91, 11303–11307 (1994).

4. Kubota H, Brinster RL. Spermatogonial stem cells. Biol Reprod 99, 52–74 (2018).

5. Lewis JP, Trobaugh FE. Hæmatopoietic Stem Cells. Nature 204, 589–590 (1964).

6. McCarrey JR. Toward a more precise and informative nomenclature describing fetal and neonatal male germ cells in rodents. Biology of Reproduction 89, 47 (2013).

7. Yang QE, Gwost I, Oatley MJ, Oatley JM. Retinoblastoma protein (RB1) controls fate determination in stem cells and progenitors of the mouse male germline. Biol Reprod 89, 113 (2013).

8. McCarrey JR. Transition of Prenatal Prospermatogonia to Postnatal Spermatogonia. In: The Biology of Mammalian Spermatogonia (ed^(eds Oatley JM, Griswold MD). Springer New York (2017).

9. Nathan C. Law MJO, Jon M. Oatley. Developmental Kinetics and Transcriptome Dynamics of Stem Cell Specification in the Spermatogenic Lineage. Nature Communications In press, (2019).

10. Russell LD, Ettlin RA, Hikim APS, Clegg ED. Histological and Histopathological Evaluation of the Testis. International Journal of Andrology 16, 83–83 (1993).

11. Lord T, Oatley JM. Regulation of Spermatogonial Stem Cell Maintenance and Self-Renewal. In: The Biology of Mammalian Spermatogonia (ed^(eds Oatley JM, Griswold MD). Springer New York (2017).

12. De Rooij DG, Russell LD. All You Wanted to Know About Spermatogonia but Were Afraid to Ask. Journal of Andrology 21, 776–798 (2000).

13. Johnson L. Spermatogenesis and Aging in the Human. Journal of Andrology 7, 331–354 (1986).

14. Allan DJ, Harmon BV, Kerr JFR, Potten CS. Perspectives on mammalian cell death. by Potten CS, Oxford University Press, London, 229–258 (1987).

15. Oakberg EF. Spermatogonial stem-cell renewal in the mouse. The Anatomical Record 169, 515–531 (1971).

16. Huckins C. The spermatogonial stem cell population in adult rats. I. Their morphology, proliferation and maturation. Anat Rec 169, 533–557 (1971).

17. Hara K, et al. Mouse Spermatogenic Stem Cells Continually Interconvert between Equipotent Singly Isolated and Syncytial States. Cell Stem Cell 14, 658–672 (2014).

18. Buaas FW, et al. Plzf is required in adult male germ cells for stem cell self-renewal. Nature Genetics 36, 647 (2004).

19. Valli H, et al. Fluorescence- and magnetic-activated cell sorting strategies to isolate and enrich human spermatogonial stem cells. Fertil Steril 102, 566–580 e567 (2014).

20. Helsel AR, Yang Q-E, Oatley MJ, Lord T, Sablitzky F, Oatley JM. ID4 levels dictate the stem cell state in mouse spermatogonia. Development 144, dev.146928-146634 (2017).

21. Hermann BP, et al. The Mammalian Spermatogenesis Single-Cell Transcriptome, from Spermatogonial Stem Cells to Spermatids. Cell Reports 25, 1650–1667.e1658 (2018).

22. Green CD, et al. A Comprehensive Roadmap of Murine Spermatogenesis Defined by Single-Cell RNA-Seq. Developmental Cell 46, 651–667.e610 (2018).

23. Guo J, et al. The adult human testis transcriptional cell atlas. Cell Research 28, 1141–1157 (2018).

24. Fayomi AP, Orwig KE. Spermatogonial stem cells and spermatogenesis in mice, monkeys and men. Stem Cell Research 29, 207–214 (2018).

25. Niedenberger BA, Busada JT, Geyer CB. Marker expression reveals heterogeneity of spermatogonia in the neonatal mouse testis. Reproduction 149, 329–338 (2015).

26. Velte EK, et al. Differential RA responsiveness directs formation of functionally-distinct spermatogonial populations at the initiation of spermatogenesis in the mouse. Development, (2019).

27. Chan F, et al. Functional and molecular features of the Id4+ germline stem cell population in mouse testes. Genes Dev 28, 1351–1362 (2014).

28. Mutoji K, et al. TSPAN8 Expression Distinguishes Spermatogonial Stem Cells in the Prepubertal Mouse Testis. Biology of Reproduction 95, 117 (2016).

29. Roadmap Epigenomics C, et al. Integrative analysis of 111 reference human epigenomes. Nature 518, 317 (2015).

30. Ram O, et al. Combinatorial Patterning of Chromatin Regulators Uncovered by Genome-wide Location Analysis in Human Cells. Cell 147, 1628–1639 (2011).

31. Zhu J, et al. Genome-wide Chromatin State Transitions Associated with Developmental and Environmental Cues. Cell 152, 642–654 (2013).

32. Gifford Casey A, et al. Transcriptional and Epigenetic Dynamics during Specification of Human Embryonic Stem Cells. Cell 153, 1149–1163 (2013).

33. Consortium EP. An integrated encyclopedia of DNA elements in the human genome. Nature 489, 57–74 (2012).

34. Kellis M, et al. Defining functional DNA elements in the human genome. Proc Natl Acad Sci U S A 111, 6131–6138 (2014).

35. Ernst J, et al. Mapping and analysis of chromatin state dynamics in nine human cell types. Nature 473, 43–49 (2011).

36. Shema E, Bernstein BE, Buenrostro JD. Single-cell and single-molecule epigenomics to uncover genome regulation at unprecedented resolution. Nature Genetics 51, 19–25 (2019).

37. Hobbs RM, Seandel M, Falciatori I, Rafii S, Pandolfi PP. Plzf Regulates Germline Progenitor Self-Renewal by Opposing mTORC1. Cell 142, 468–479 (2010).

38. Lee J, et al. Akt mediates self-renewal division of mouse spermatogonial stem cells. Development 134, 1853–1859 (2007).

39. Aibar S, Fontanillo C, Droste C, De Las Rivas J. Functional Gene Networks: R/Bioc package to generate and analyse gene networks derived from functional enrichment and clustering. Bioinformatics 31, 1686–1688 (2015).

40. Elsässer SJ, Noh K-M, Diaz N, Allis CD, Banaszynski LA. Histone H3.3 is required for endogenous retroviral element silencing in embryonic stem cells. Nature 522, 240–244 (2015).

41. Calo E, Wysocka J. Modification of Enhancer Chromatin: What, How, and Why? Molecular Cell 49, 825–837 (2013).

42. Shen Y, et al. A map of the cis-regulatory sequences in the mouse genome. Nature 488, 116–120 (2012).

43. Hammoud SS, Low DHP, Yi C, Carrell DT, Guccione E, Cairns BR. Chromatin and Transcription Transitions of Mammalian Adult Germline Stem Cells and Spermatogenesis. Cell Stem Cell 15, 239–253 (2014).

44. Oatley JM, Oatley MJ, Avarbock MR, Tobias JW, Brinster RL. Colony stimulating factor 1 is an extrinsic stimulator of mouse spermatogonial stem cell self-renewal. Development 136, 1191–1199 (2009).

45. Stadler MB, et al. DNA-binding factors shape the mouse methylome at distal regulatory regions. Nature 480, 490 (2011).

46. Wang H, et al. Widespread plasticity in CTCF occupancy linked to DNA methylation. Genome Res 22, 1680–1688 (2012).

47. La HM, et al. Identification of dynamic undifferentiated cell states within the male germline. Nature Communications 9, (2018).

48. Matzuk MM, Lamb DJ. The biology of infertility: research advances and clinical challenges. Nature Medicine 14, 1197–1213 (2008).

49. Jelinic P, Stehle J-C, Shaw P. The testis-specific factor CTCFL cooperates with the protein methyltransferase PRMT7 in H19 imprinting control region methylation. PLOS Biology 4, e355 (2006).

50. Zhang T, Oatley J, Bardwell VJ, Zarkower D. DMRT1 Is Required for Mouse Spermatogonial Stem Cell Maintenance and Replenishment. PLOS Genetics 12, (2016).

51. Zhang T, Murphy MW, Gearhart MD, Bardwell VJ, Zarkower D. The mammalian Doublesex homolog DMRT6 coordinates the transition between mitotic and meiotic developmental programs during spermatogenesis. Development 141, 3662–3671 (2014).

52. Ran LL, et al. FOXF1 Defines the Core-Regulatory Circuitry in Gastrointestinal Stromal Tumor. Cancer Discov 8, 234–251 (2018).

53. Gemenetzidis E, Elena-Costea D, Parkinson EK, Waseem A, Wan H, Teh MT. Induction of human epithelial stem/progenitor expansion by FOXM1. Cancer Res 70, 9515–9526 (2010).

54. Besharat ZM, et al. Foxm1 controls a pro-stemness microRNA network in neural stem cells. Sci Rep 8, 3523 (2018).

55. Goertz MJ, Wu Z, Gallardo TD, Hamra KF, Castrillon DH. Foxo1 is required in mouse spermatogonial stem cells for their maintenance and the initiation of spermatogenesis. Journal of Clinical Investigation 121, 3456–3466 (2011).

56. Guo J, et al. Chromatin and Single-Cell RNA-Seq Profiling Reveal Dynamic Signaling and Metabolic Transitions during Human Spermatogonial Stem Cell Development. Cell Stem Cell 21, 533–546.e536 (2017).

57. Leishman E, et al. Foxp1 maintains hair follicle stem cell quiescence through regulation of Fgf18. Development 140, 3809–3818 (2013).

58. Li HJ, et al. FOXP1 controls mesenchymal stem cell commitment and senescence during skeletal aging. Journal of Clinical Investigation 127, 1241–1253 (2017).

59. Naudin C, et al. PUMILIO/FOXP1 signaling drives expansion of hematopoietic stem/progenitor and leukemia cells. Blood 129, 2493–2506 (2017).

60. Gabut M, et al. An Alternative Splicing Switch Regulates Embryonic Stem Cell Pluripotency and Reprogramming. Cell 147, 132–146 (2011).

61. Sarkar A, Hochedlinger K. The Sox Family of Transcription Factors: Versatile Regulators of Stem and Progenitor Cell Fate. Cell Stem Cell 12, 15–30 (2013).

62. Raverot G, Weiss J, Park SY, Hurley L, Jameson JL. Sox3 expression in undifferentiated spermatogonia is required for the progression of spermatogenesis. Developmental Biology 283, 215–225 (2005).

63. Ernst J, Kellis M. Chromatin-state discovery and genome annotation with ChromHMM. Nat Protoc 12, (2017).

64. Murphy MW, et al. An ancient protein-DNA interaction underlying metazoan sex determination. Nat Struct Mol Biol 22, 442–U426 (2015).

65. Rivero-Hinojosa S, Kang S, Lobanenkov VV, Zentner GE. Testis-specific transcriptional regulators selectively occupy BORIS-bound CTCF target regions in mouse male germ cells. Sci Rep 7, 41279 (2017).

66. Ren G, et al. CTCF-Mediated Enhancer-Promoter Interaction Is a Critical Regulator of Cell-to-Cell Variation of Gene Expression. Mol Cell 67, 1049–1058 e1046 (2017).

67. Derrien T, et al. The GENCODE v7 catalog of human long noncoding RNAs: Analysis of their gene structure, evolution, and expression. Genome Research, 1775–1789 (2012).

68. Zacher B, Michel M, Schwalb B, Cramer P, Tresch A, Gagneur J. Accurate Promoter and Enhancer Identification in 127 ENCODE and Roadmap Epigenomics Cell Types and Tissues by GenoSTAN. Plos One 12, e0169249 (2017).

69. Lesch BJ, Dokshin GA, Young RA, McCarrey JR, Page DC. A set of genes critical to development is epigenetically poised in mouse germ cells from fetal stages through completion of meiosis. P Natl Acad Sci USA 110, 16061–16066 (2013).

70. Zentner GE, of Chemistry S-PC. The chromatin fingerprint of gene enhancer elements. Journal of Biological Chemistry, (2012).

71. Nakagawa T, Sharma M, Nabeshima Y-I, Braun RE, Yoshida S. Functional Hierarchy and Reversibility Within the Murine Spermatogenic Stem Cell Compartment. Science (New York, NY) 328, 62–67 (2010).

72. Murphey P, McLean DJ, McMahan CA, Walter CA, McCarrey JR. Enhanced genetic integrity in mouse germ cells. Biology of Reproduction 88, 6 (2013).

73. Brind’Amour J, Liu S, Hudson M, Chen C, Karimi MM, Lorincz MC. An ultra-low-input native ChIP-seq protocol for genome-wide profiling of rare cell populations. Nature Communications 6, 6033 (2015).

74. Taiwo O, et al. Methylome analysis using MeDIP-seq with low DNA concentrations. Nat Protoc 7, 617–636 (2012).

75. Liao Y, Smyth GK, Shi W. The R package Rsubread is easier, faster, cheaper and better for alignment and quantification of RNA sequencing reads. Nucleic Acids Res, (2019).

76. Gaidatzis D, Lerch A, Hahne F, Stadler MB. QuasR: quantification and annotation of short reads in R. Bioinformatics (Oxford, England) 31, 1130–1132 (2015).

77. Liao Y, Smyth GK, Shi W. featureCounts: an efficient general purpose program for assigning sequence reads to genomic features. Bioinformatics 30, 923–930 (2014).

78. Love MI, Huber W, Anders S. Moderated estimation of fold change and dispersion for RNA-seq data with DESeq2. Genome biology 15, 550 (2014).

79. Lun A, Smyth GK. csaw: a Bioconductor package for differential binding analysis of ChIP-seq data using sliding windows. Nucleic Acids Research 44, (2016).

80. Scrucca L, Fop M, Murphy TB, Raftery AE. mclust 5: Clustering, Classification and Density Estimation Using Gaussian Finite Mixture Models. The R journal 8, 289–317 (2016).

81. Ramirez F, et al. deepTools2: a next generation web server for deep-sequencing data analysis. Nucleic Acids Res 44, W160–165 (2016).

82. Shen L, Shao N, Liu X, Nestler E. ngs.plot: Quick mining and visualization of next-generation sequencing data by integrating genomic databases. BMC Genomics 15, 284 (2014).

83. Lienhard M, Grimm C, Morkel M, Herwig R, Chavez L. MEDIPS: genome-wide differential coverage analysis of sequencing data derived from DNA enrichment experiments. Bioinformatics (Oxford, England) 30, 284–286 (2013).

84. Zhang Y, et al. Model-based Analysis of ChIP-Seq (MACS). Genome Biology 9, R137 (2008).

85. Yu G, Wang LG, He QY. ChIPseeker: an R/Bioconductor package for ChIP peak annotation, comparison and visualization. Bioinformatics 31, 2382–2383 (2015).

86. Yu G. clusterProfiler: An universal enrichment tool for functional and comparative study. bioRxiv, 256784 (2018).

87. McLean CY, et al. GREAT improves functional interpretation of cis-regulatory regions. Nature Biotechnology 28, 495–501 (2010).

88. Yue F, et al. A comparative encyclopedia of DNA elements in the mouse genome. Nature 515, 355 (2014).

89. Ramirez F, et al. High-resolution TADs reveal DNA sequences underlying genome organization in flies. Nature Communications 9, 189 (2018).

90. Ye J, Coulouris G, Zaretskaya I, Cutcutache I, Rozen S, Madden TL. Primer-BLAST: A tool to design target-specific primers for polymerase chain reaction. BMC Bioinformatics 13, 134 (2012).

91. Serra ND, Velte EK, Niedenberger BA, Kirsanov O, Geyer CB. Cell-autonomous requirement for mammalian target of rapamycin (Mtor) in spermatogonial proliferation and differentiation in the mouse†. Biology of Reproduction 96, 816–828 (2017).

