## Supplementary Figures for "Unique Epigenetic Programming Distinguishes Regenerative Spermatogonial Stem Cells in the Developing Mouse Testis"

#### Figure S1

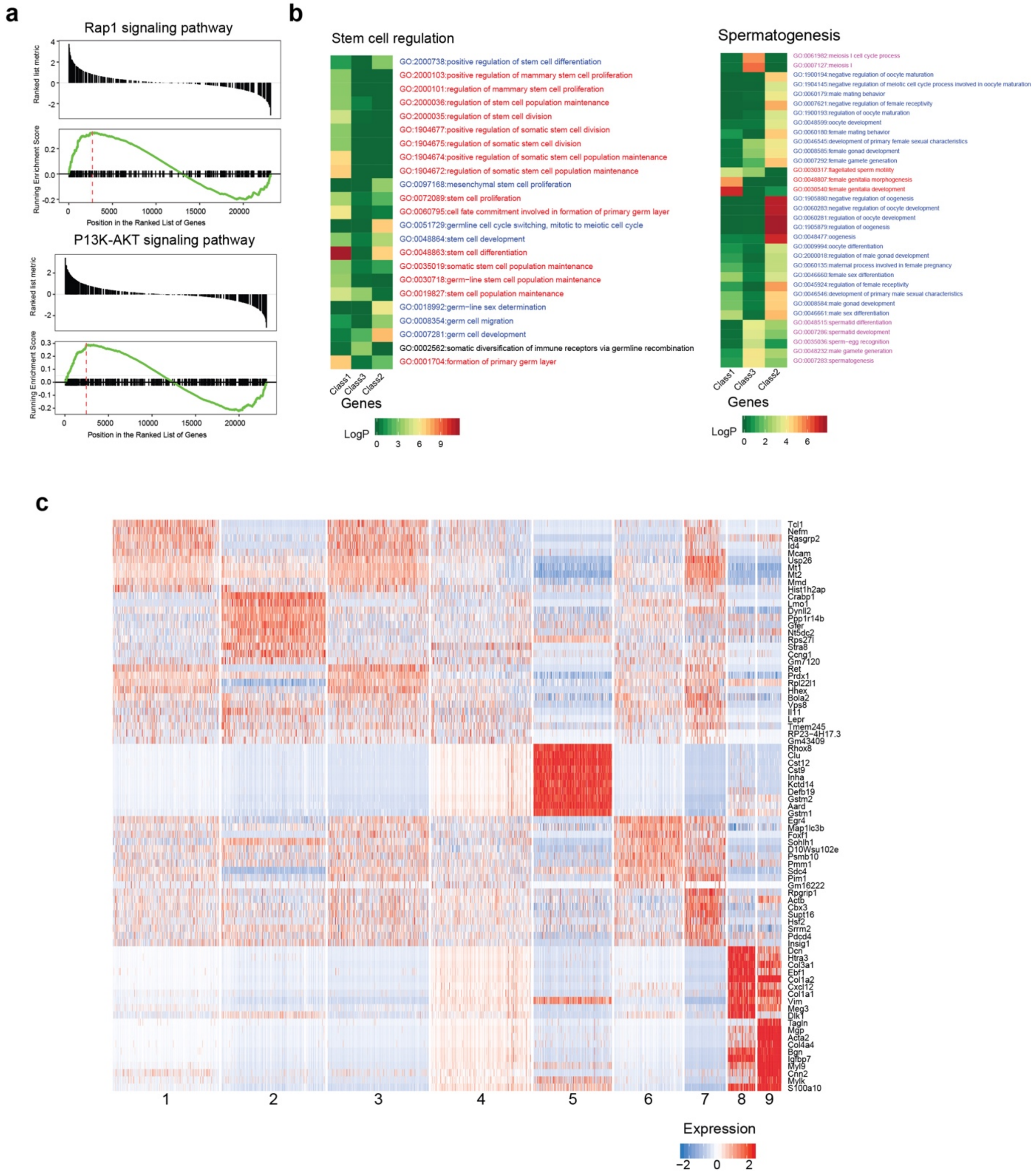


**Fig. S1: Bulk RNA-seq and scRNA-seq analysis of gene expression in ID4-eGFP$^{Bright}$ and ID4-eGFP$^{Dim}$ cells. a** Gene Set Enrichment Analysis (GSEA) of KEGG pathways detected among genes similarly expressed in ID4-eGFP$^{Bright}$ and ID4-eGFP$^{Dim}$ spermatogonia. The Rap1 signaling pathway and P13K-AKT signaling pathway signaling pathways are utilized to a similar extent in both ID4-eGFP$^{Bright}$ and ID4-eGFP$^{Dim}$ spermatogonia. **b** Heatmaps of gene ontology (Stem cell regulation and Spermatogenesis enrichment analysis of Class 1, Class 2 and Class 3 genes expressed in ID4-eGFP$^{Bright}$ and/or ID4-eGFP$^{Dim}$ spermatogonia. **c** Heatmap display of expression of the top 10 ranked marker genes included within each cell cluster shown in **Fig. 1b**.

### Figure S2

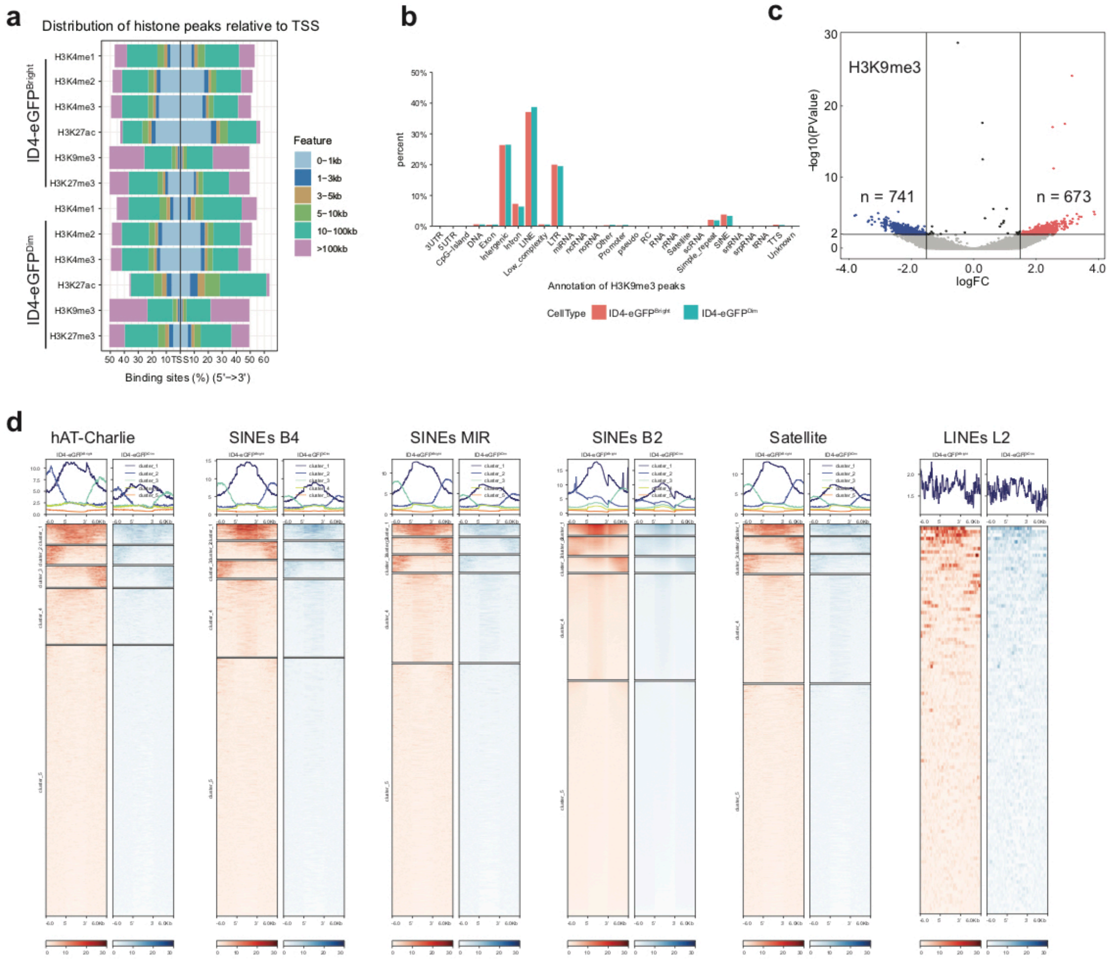


**e**

| | Enriched GO Term | P Value |
|---|---|---|
| Up in ID4-eGFP^Bright | stem cell population maitennance | 4.79e-04 |
| | formation of primary germ layer | 6.72e-03 |
| | negative regulation of stem cell differentiaon | 1.56e-02 |
| | regulation of stem cell diffrention | 2.32e-02 |
| | stem cell differentiation | 3.87e-02 |
| | regulation of stem cell population maitenance | 4.82e-02 |
| | asymetric stem cell division | 8.53e-02 |
| Up in ID4-eGFP^Dim | meiosis I | 1.13e-03 |
| | male meiosis | 6.01e-03 |
| | germ cell development | 9.28e-03 |
| | oogenesis | 1.74e-02 |
| | oocyte differentiation | 3.74e-02 |
| | female gamete generation | 4.01e-02 |
| | male gamete generation | 6.49e-02 |
| | spermatogenesis | 6.51e-02 |

Diffterential H3K27ac sites at promoters

**f**

| | Enriched GO Term | P Value |
|---|---|---|
| Up in ID4-eGFP^Bright | female sex differentiation | 7.74e-04 |
| | male sex differentiation | 9.42e-04 |
| | male genitalia development | 1.64e-03 |
| | female gonad development | 1.95e-03 |
| | neuronal stem cell population maintenance | 1.99e-03 |
| | regulation of male gonad development | 6.72e-03 |
| | negative regulation of female gonad development | 7.86e-03 |
| Up in ID4-eGFP^Dim | formation of primary germ layer | 1.02e-07 |
| | stem cell differentiation | 3.62e-06 |
| | positive regulation of stem cell differentiation | 6.68e-04 |
| | stem cell proliferation | 1.44e-03 |
| | male sex differentiation | 2.57e-03 |
| | male gonad development | 6.68e-03 |
| | mesenchymal stem cell differentiation | 1.33e-02 |
| | stem cell population maintenance | 1.78e-02 |

Diffterential H3K27me3 sites at promoters

**Fig. S2: Genomic features of histone modifications.** **a** Patterns of histone modifications relative to transcriptional start sites. **b** Patterns of histone modifications relative to specific genomic features. **c** Site-specific differences in enrichment of the H3K9me3 modificaton distinguishing mouse ID4-eGFP$^{Bright}$ and ID4-eGFP$^{Dim}$ spermatogonia. Red dots denote significantly enriched reads in ID4-eGFP$^{Bright}$ cells and blue dots denote significantly enriched reads in ID4-eGFP$^{Dim}$ cells. **d** Heatmaps showing relative enrichment of the H3K9me3 modification within different types of repetitive elements. **e** Gene Ontology analysis of genes showing spermatogonial-subtype specific differences in enrichment of the H3K27ac modification at promoters. **f** Gene Ontology analysis of genes showing spermatogonial-subtype specific differences in enrichment of the H3K27me3 modification at promoters.

#### Figure S3

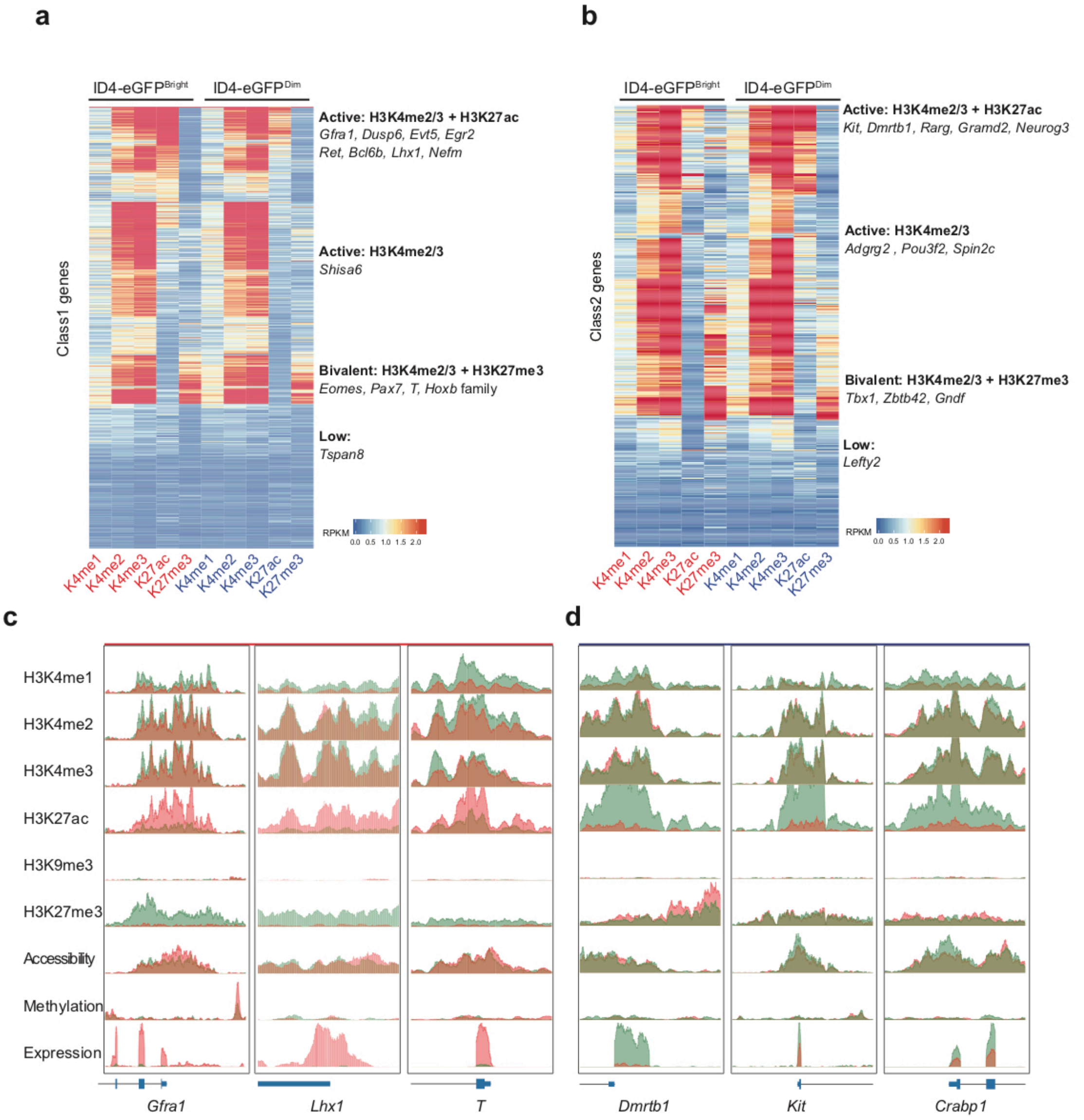


**Fig. S3: Coordinated histone modifications associated with differentially expressed genes.** Combinations of histone modifications found at active, bivalent or low level promoters of Class 1 (**a**) and Class 2 (**b**) genes in ID4-eGFP$^{Bright}$ and ID4-eGFP$^{Dim}$ spermatogonia. **c, d** Genomic browser screenshots of promoters of Class 1 (**c**) and Class 2 (**d**) genes showing relative enrichment of six different histone modifications (H3K4me1-3, H3K27ac, H3K9me3, H3K27me3) detected by ChIP-seq, chromatin accessibility detected by ATAC-seq, and DNA methylation detected by MeDIP-seq differentially expressed in ID4-eGFP$^{Bright}$ and ID4-eGFP$^{Dim}$ spermatogonia. Coral color = enrichment in ID4-eGFP$^{Bright}$ spermatogonia, green color = enrichment in ID4-eGFP$^{Dim}$ spermatogonia, dark red or brown = overlapping enrichment in both ID4-eGFP$^{Bright}$ and D4-eGFP$^{Dim}$ spermatogonia.

#### Figure S4

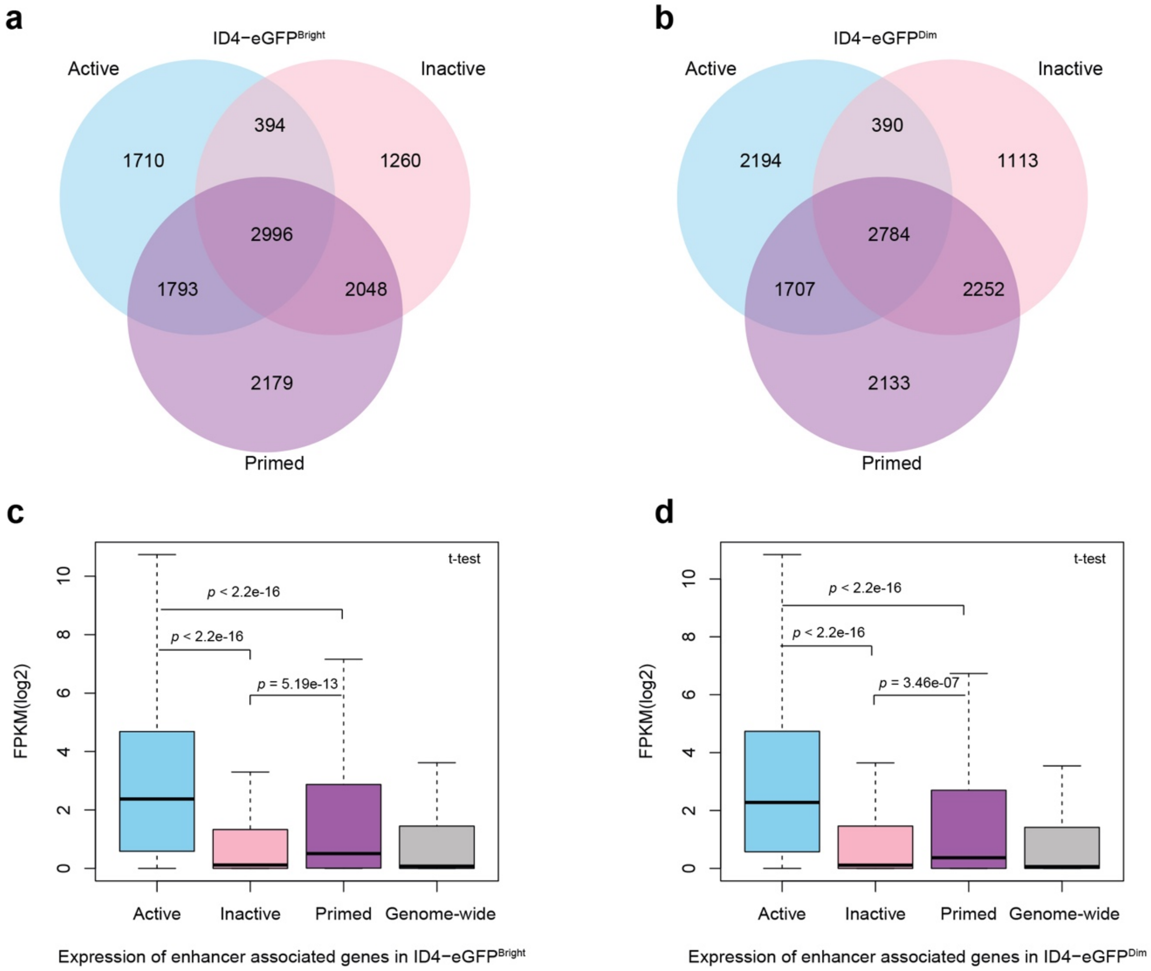


**Fig. S4: Differential expression of enhancer-associated genes. a, b** Enhancer-associated genes in ID4-eGFP$^{Bright}$ and ID4-eGFP$^{Dim}$spermatogonia. Many genes are associated with more than one enhancer, and those enhancers can be in different states. **c, d** GREAT GO analysis of expression levels of genes associated with different types of enhancers in ID4-eGFP$^{Bright}$ and ID4-eGFP$^{Dim}$ spermatogonia. Genes associated with active enhancers were expressed at significantly higher levels than those associated with either inactive or primed enhancers, or than all genes combined in each spermatogonial subtype.

#### Figure S5

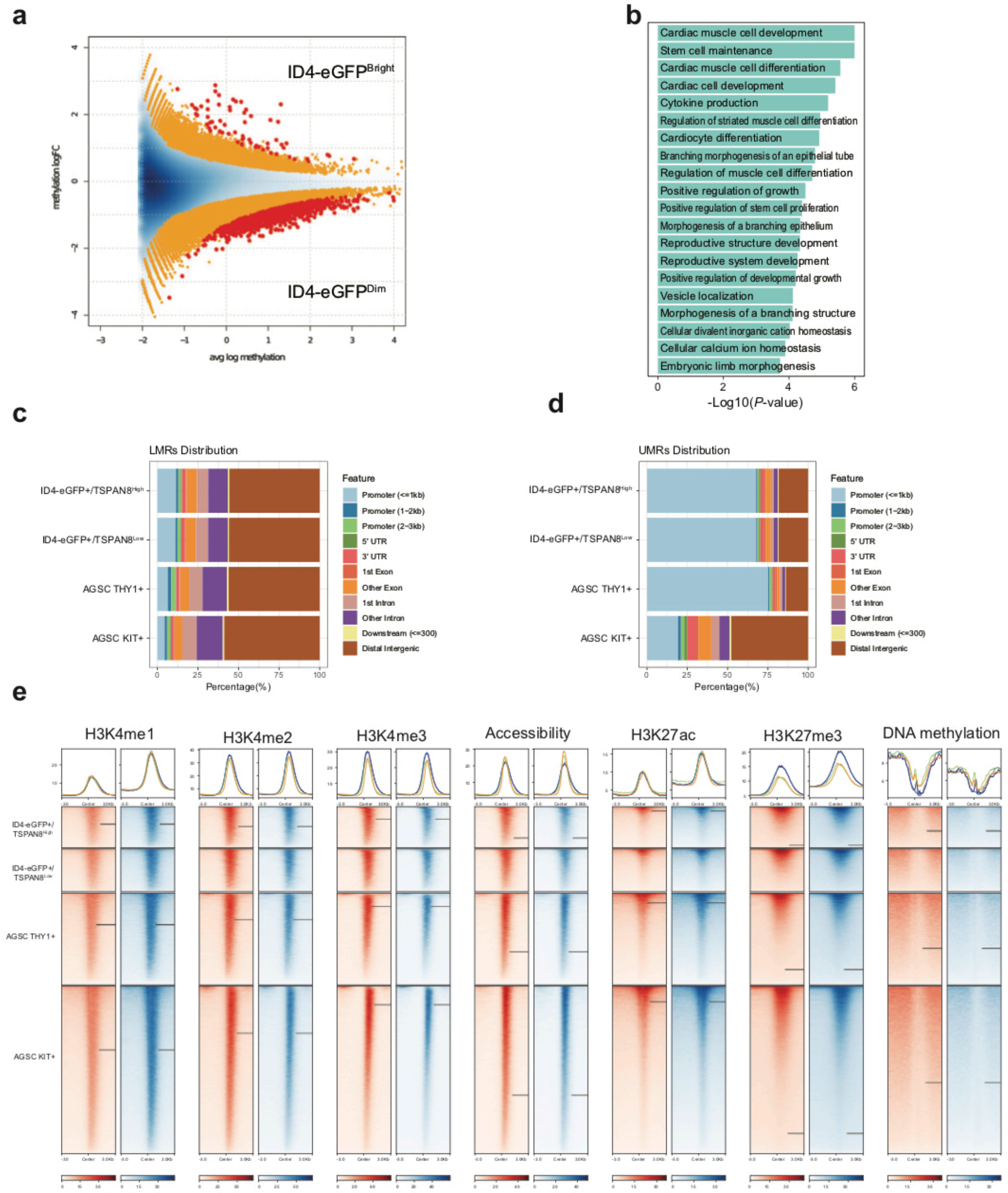


**Fig. S5: Characteristics of differential DNA methylation patterns in SSC-enriched ID4-eGFP$^{Bright}$ and progenitor-enriched ID4-eGFP$^{Dim}$ spermatogonia. a** Volcanoe plot showing significantly differentially methylated regions (DMRs) or sites depicted as red dots (FDR < 0.1) in ID4-eGFP$^{Bright}$ and ID4-eGFP$^{Dim}$ spermatogonia. Each red dot represents an individual CpG dinucleotide at which the level of DNA methylation is statistically significantly different in one spermatogonial subtype compared to the other. CpGs represented in orange and blue regions did not show significantly different methylation levels. **b** GREAT-GO gene ontology analysis showed that spermatogonial-subtype specific DMRs were associated with genes involved in critical functions including stem cell maintenance, proliferation, morphogenesis and differentiation. **c,d** Genomic distribution of LMRs (**c**) and UMRs (**d**) shows that most LMRs occur in distal intergenic regions whereas most UMRs occur in gene promoter regions. **e** ChIP-seq analysis showed that LMRs align with sites of concentrated enrichment of five different histone modifications (H3K4me1-3, H3K27ac, H3K27me3) and peaks of chromatin accessibility in all four spermatogonial subpopulations, suggesting LMRs demarcate sites of intergenic regulatory elements.

**Figure S6**

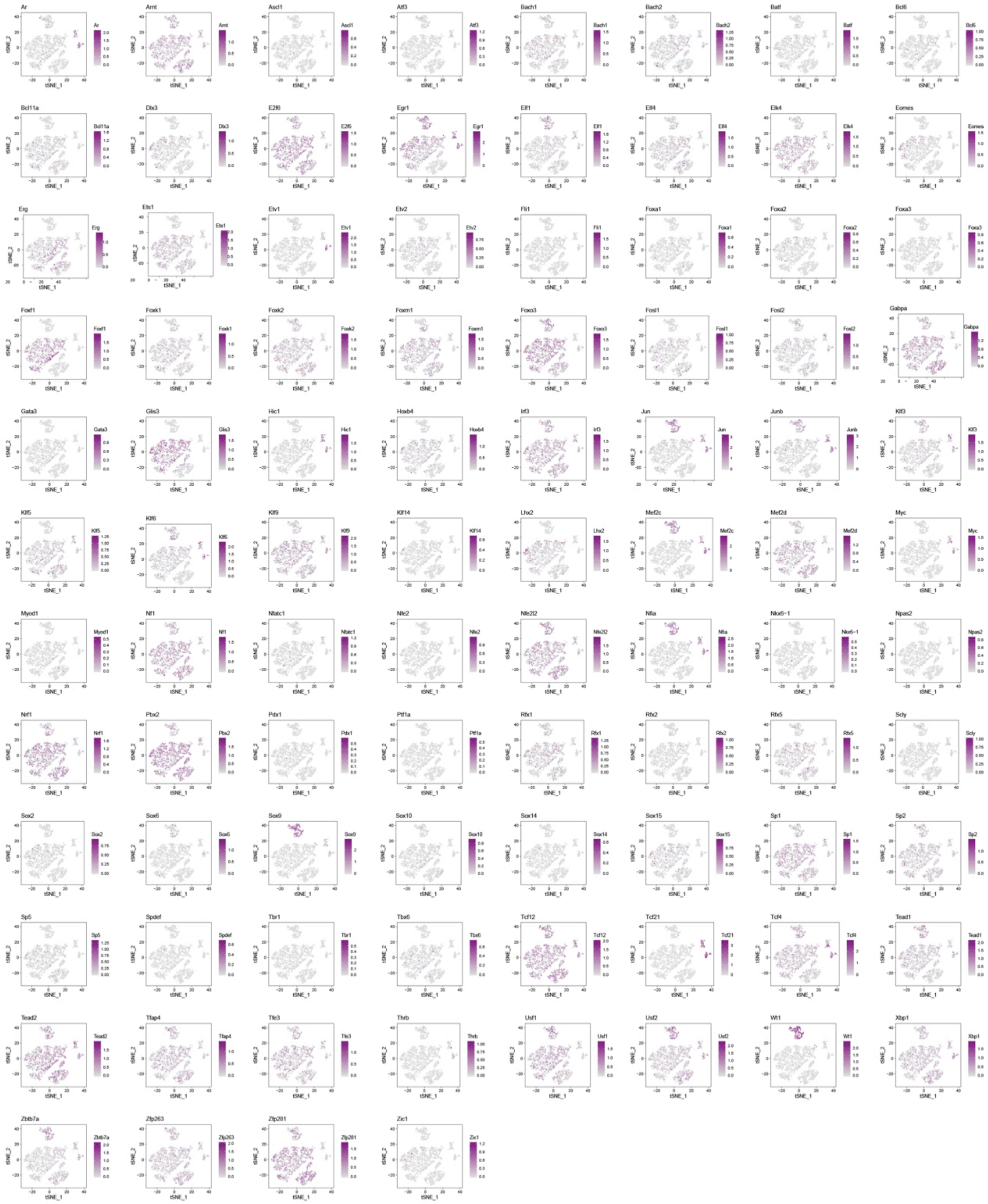

**Fig. S6: Cell type/subtype-specific expression patterns of mRNAs encoding candidate regulators of epigenetic programming in ID4-eGFP$^{Bright}$ and ID4-eGFP$^{Dim}$ spermatogonia.** scRNA-seq data showing expression of mRNAs encoding transcription factors predicted by motif enrichment analysis to bind to enhancer regions in ID4-eGFP$^{Bright}$ and/or ID4-eGFP$^{Dim}$ spermatogonia.

#### Figure S7

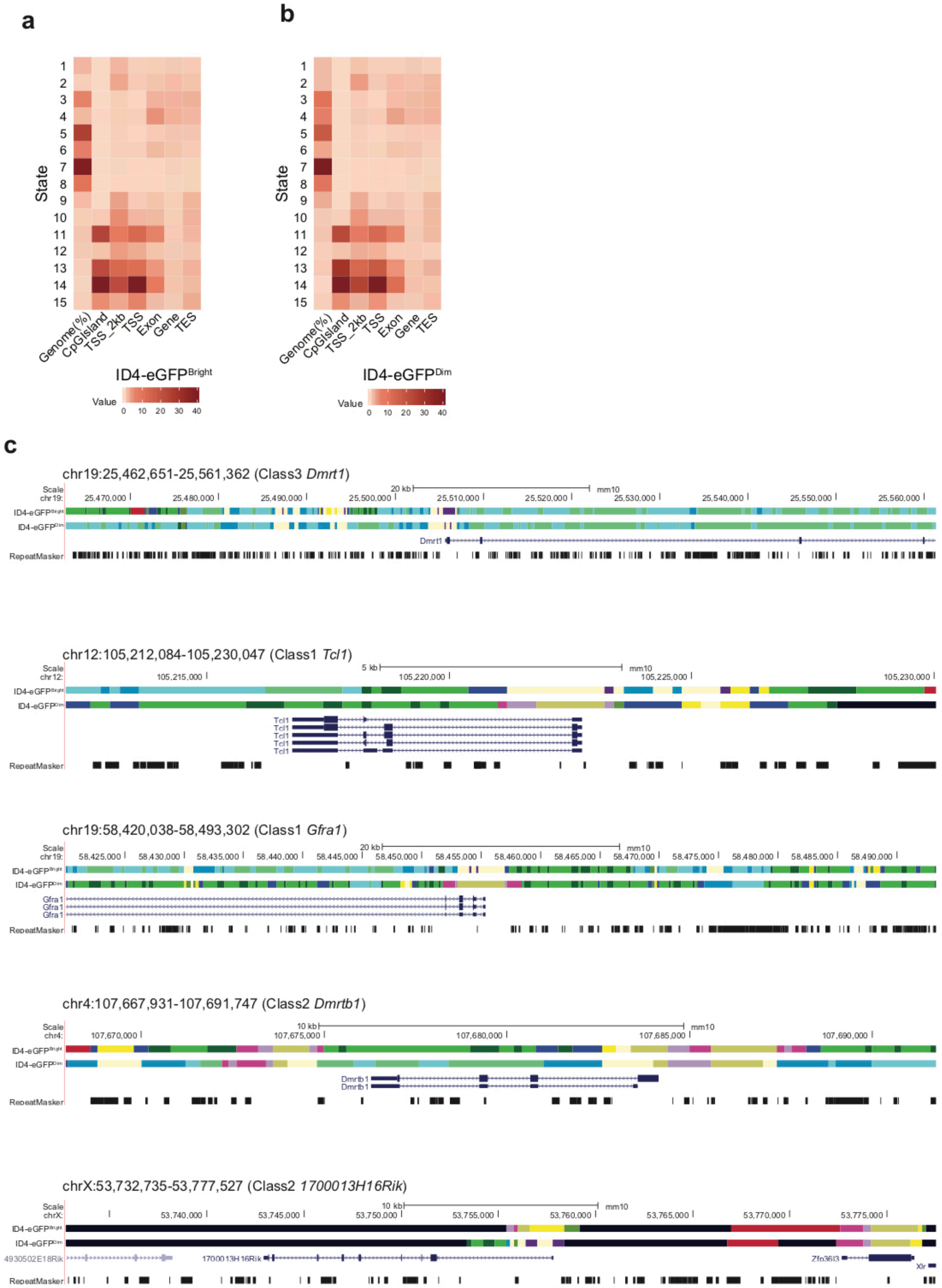


**Fig. S7: Differential chromatin states are associated with spermatogonial subtype-specific differential gene expression. a** The relative genome-wide distribution of each of the 15 chromatin states as a function of specific genomic features in ID4-eGFP$^{Bright}$ (**a**) and ID4-eGFP$^{Dim}$ (**b**) spermatogonial subtypes. **c** Browser views of ChromHMM genome annotations representative of the 15 different chromatin states described in **a** and **b**. in one similarly expressed Class 3 gene (*Dmrt1*), two differentially expressed Class 1 genes (*Tcl1* & *Gfra1*), and two differentially expressed Class 2 genes (*Dmrtb1*). Color coding of chromatin states is as shown in **Fig. 7a**.
